# Subregions of the infant hippocampal-entorhinal system are linked to distinct cortical networks in the first two years of human life

**DOI:** 10.64898/2026.10.01.755949

**Authors:** Daniel Reznik, Cheslie C. Klein, Shir Filo, Menno P. Witter, Charlotte Grosse Wiesmann

## Abstract

Episodic memory critically depends on the hippocampal-entorhinal system. Despite a rapid developmental change in human episodic memory during the first two years of life, little is known about the developmental trajectory of the hippocampal-entorhinal circuitry during this critical time period. In the current study, we collected high-quality precision fMRI data from five typically developing infants over the time period between 1 and 2 years of life and examined the cortical networks associated with subregions of the hippocampal–entorhinal system. We show that already at around 1 year of age, subdivisions of the canonical default network (DN) are dissociated at the neocortex but are differently associated with the hippocampal-entorhinal system, with tentatively differential developmental trajectories. While the DN-A, associated with episodic simulation, coupled with the anterior hippocampus and the medial part of the entorhinal cortex, the DN-B, associated with social cognition, showed weaker connectivity with the hippocampal-entorhinal system. The parietal memory network, in turn, coupled with the posterior hippocampus and the lateral part of the entorhinal cortex. Exploratory longitudinal results suggest that associations of the DN-B with the hippocampus and the intermediate part of the entorhinal cortex increase towards the end of the second postnatal year, and the associations of the parietal memory network become more topographically restricted to the posterior hippocampus. These networks and their differential maturation may provide the basis for important developments in episodic memory and social cognition in the second postnatal year and suggest that potentially unique properties of human hippocampal-entorhinal circuitry may emerge latest in development.

## Introduction

Episodic memory - conscious specific memory for own life events – critically depends on the brain regions comprising the medial temporal lobe (MTL), particularly the hippocampus and the entorhinal cortex^1–3^. Although the hippocampus is involved in human mnemonic processing already from around one year of age^4^, it is not typically before the end of the second postnatal year that children show robust evidence of recalling specific and context-dependent elements of past experiences^5–8^. While the development of episodic memory in humans has been proposed to be associated with the accelerated structural maturation of the hippocampus in the first two year of life^4,9–11^, remarkably little is known about the maturation of the human hippocampal-entorhinal circuitry and how it becomes integrated with distributed cortical processing in this critical period.

In contrast to humans, animal anatomical studies provided critical insights into the maturation process of the hippocampal-entorhinal circuitry indicating that distinct components of this circuitry mature at different developmental stages. For example, posterior subdivisions of the primate entorhinal cortex mature earlier than its anterior subdivisions^12,13^. Since the entorhinal cortex provides most of the cortical input to the hippocampus^2,14^, the modular developmental pattern of the animal entorhinal cortex suggests that the emergence of adult-like hippocampal-dependent functions occurs via a gradual maturation of distinct subcomponents of the circuitry associating the entorhinal cortex with the hippocampus and with distributed neocortical regions.

Recent methodological advances in functional magnetic resonance imaging (fMRI) have made it possible to investigate the hippocampal-entorhinal circuitry in the adult human brain at the individual-subject level with unprecedented anatomical detail. By moving away from the classical group-average approach, individualized neuroimaging has demonstrated that distinct parts of the hippocampus and entorhinal cortex are associated with at least three distributed cortical networks^15–20^ – two subdivisions of the canonical default network^21–23^, Default Network A (DN-A) and Default Network B (DN-B)^24–27^, and the Parietal Memory Network, also referred to as the Salience Network (PMN/SAL)^18,28–30^. Specifically, the most medial part of the human entorhinal cortex, closest to the hippocampus, is associated with the DN-A^15^. The more intermediately positioned part of the entorhinal cortex is associated with the DN-B^15,16^ (see Ref^17^ for an independent replication), and its most lateral part, closest to the perirhinal cortex, is associated with the PMN^16,31^. Importantly, these networks are dissociable not only in terms of their anatomical organization, but also in their functional properties. In adults, the DN-A is selectively implicated in episodic simulation, the DN-B in social cognition and mentalizing^26,27^, and the PMN/SAL in recognition memory and salience detection^20,32^. Finally, these distinct parts of the human entorhinal cortex and their associated cortical networks are linked with distinct parts of the hippocampal long axis^16^, together forming a clearly organized and phylogenetically conserved topographical architecture supporting episodic memory and other hippocampus-dependent cognitive functions in the mammalian lineage^33–35^.

Despite these recent breakthroughs in mapping the organization of the hippocampal-entorhinal circuitry in human adults—and in striking contrast to our knowledge of its development in animals—how this circuitry matures into its adult form and how associations of its distinct parts with distinct cortical networks develop during human infancy remain unknown. This gap in knowledge has been exceptionally difficult to address due to long-standing challenges of acquiring dense high-quality fMRI data from individual infants^36^ as well as due to substantial blurring of fine-grained MTL anatomy in group-averaged analyses^27^.

In this study, we addressed these challenges by leveraging within-individual precision mapping with extensive high-quality fMRI data to characterize the distributed cortical circuitry associated with the hippocampal-entorhinal system in five human infants between 10 and 16 months of age. To trace developmental change across the functionally critical window between 1 and 2 years of age, we further acquired longitudinal precision fMRI data from two of these infants at approximately 2 years of age. Using two independent datasets for network discovery and validation within each individual, we show that, at the neocortical level, the networks DN-A and DN-B are already fully dissociated and follow an adult-like topography by 10 months of age. While the DN-A and PMN/SAL show mature connectivity with the MTL at this age, the connectivity of the DN-B with the MTL is significantly weaker. Our exploratory longitudinal findings suggest that connectivity of the DN-B with the infant hippocampal system may develop later, between 1.5 and 2 years, and that the association of the PMN/SAL with the posterior hippocampus, in turn, may become increasingly topographically restricted over this same developmental interval. Together, these findings suggest distinct developmental trajectories of hippocampal-entorhinal networks in the first two years of human life.

## Results

### Data quality

Spontaneous brain activity was recorded from five infants during natural sleep using fMRI (Table 1). Since head movement during data acquisition dramatically affects functional connectivity estimates and introduces spurious correlations^37,38^, we first excluded all BOLD runs with absolute maximum head displacement greater than 2.2 mm from the analyses. Consequently, maximum head displacement across all remaining BOLD runs was 2.12 mm for participant 1 (P1), 1.37 mm for P2, 1.3 mm for P3, 1.74 mm for P4, and 1.67 mm for P5. Next, we examined whole-brain signal quality in each infant by calculating tSNR values for each individual voxel. Signal quality was excellent at the whole-brain level (Figure 1, upper row), and importantly, also in the medial temporal lobe (Figure 1, bottom row; see Figure S1 for data from other participants). Notably, signal quality in the infant entorhinal cortex was better than typical signal quality in individual adult subjects^39^. While tSNR values were still lower in the most ventral compared with the dorsal parts of the infant entorhinal cortex, they remained within an acceptable range.

**Figure 1.**
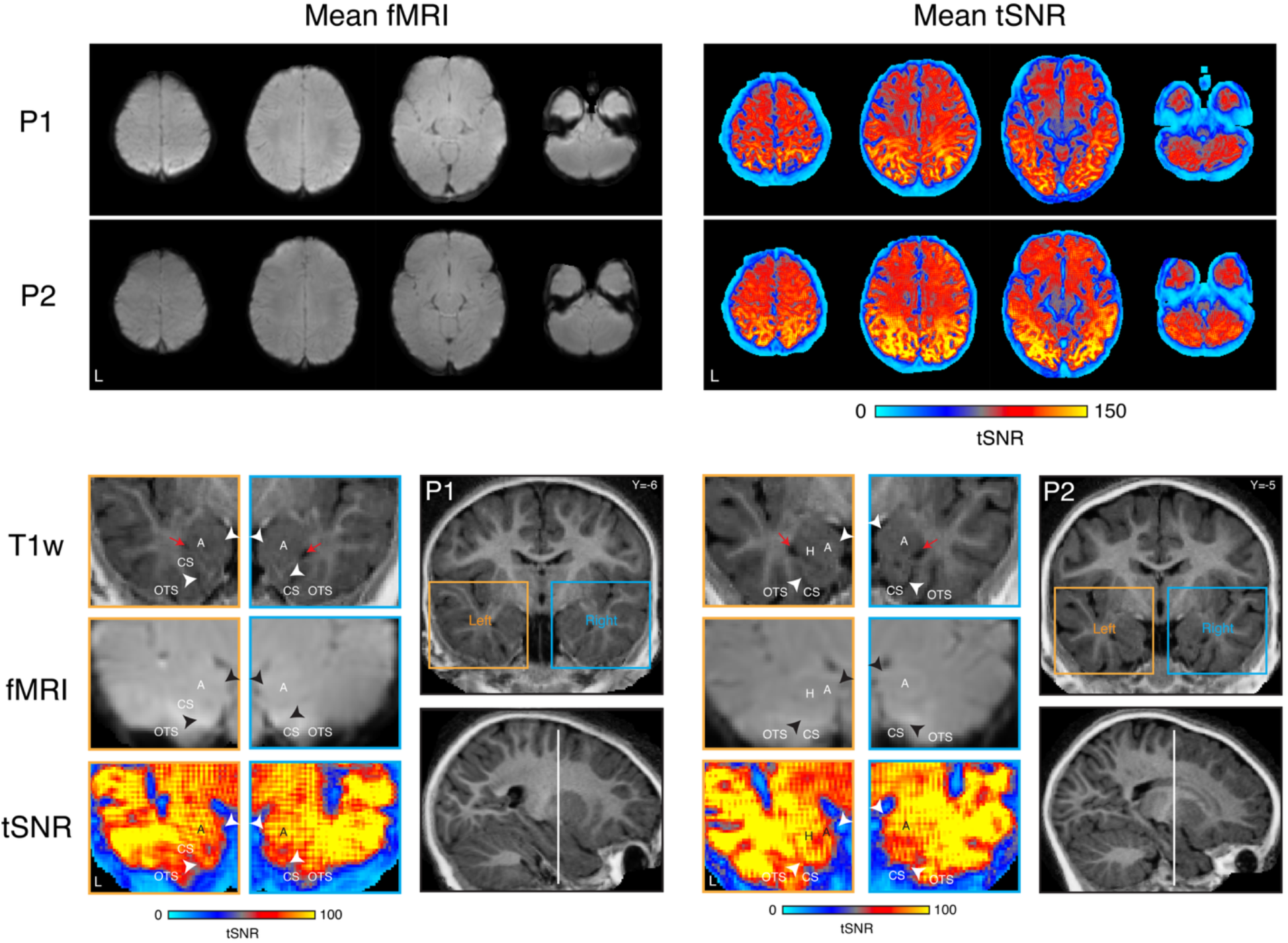
Infant fMRI data quality. Upper row – whole-brain data quality. Axial slices from P1 and P2 show mean functional (fMRI; left) and mean tSNR (right) data. High coverage and high tSNR were obtained in all regions of the brain, particularly in the medial temporal lobe. Bottom row – data quality in the infant medial temporal lobe. Coronal slices showing the hippocampus and entorhinal cortex at the level of amygdala presented using three signal modalities in P1 (left) and P2 (right) – anatomical data (T1w), functional data (fMRI) and tSNR data (tSNR). The putative medial-lateral extent of the entorhinal cortex is marked with white and black arrowheads. As can be seen from the corresponding images, brain coverage and tSNR were excellent for the entire medial-lateral extent of the infant entorhinal cortex. See Figure S1 for data from other participants. A – amygdala; cs – collateral sulcus; H – hippocampus; OTS – occipital-temporal sulcus.

**Table 1.** Infant sample data. We sampled 5 infants aged from 10 to 16 months at time point 1. About 8 months later, at time point 2, we analyzed additional data from 2 of the original 5 infants at nearly 2 years of age. Participant ID was ranked based on the mean age across the scanning sessions at time point 1. Here as well as throughout the manuscript, infants’ age was transformed to a digital number representation and is measured by months following birth. All infants sampled for this study exhibited normal general development, measured with the Bayley Scales of Infant and Toddler Development III^40^ (measured at time point 1; column 4) whose raw scores followed infants’ age. For each infant, we aimed to collect about one hour of good quality spontaneous data recorded using fMRI during natural sleep. Columns 5 and 7 shows the amount of data in minutes retained at time point 1 and 2, respectively, after removing runs with excessive head movement.

| **Participant ID** | **Sex** | **Age**<br>***Time 1*** | **Bayley**<br>**Score** | **Amount of data (min)**<br>***Time 1*** | **Age**<br>***Time 2*** | **Amount of data (min)**<br>***Time 2*** |
| --- | --- | --- | --- | --- | --- | --- |
| P1 | F | 16.1 | 76 | 72 | 23.9 | 24 |
| P2 | F | 15.5 | 77 | 60 | 24.0 | 48 |
| P3 | F | 12.6 | 64 | 60 | - | - |
| P4 | F | 12.3 | 65 | 54 | - | - |
| P5 | M | 10.8 | 56 | 72 | - | - |

### Discovery of infant distributed cortical networks and their associations with the hippocampal-entorhinal system

We aimed to examine whether the cortical networks that are associated with distinct components of the adult hippocampal-entorhinal system are already present in human infants between 10 and 16 months of age. Furthermore, our aim was to investigate how these networks associate with the infant hippocampal-entorhinal system. Using discovery datasets (half of the data for each infant), we placed seeds in the posterior midline cortex (cingulate and retrosplenial cortex; see Methods for more details on initial seed selection and placement) to identify potential candidates for the cortical networks DN-A, DN-B and PMN/SAL in each infant. We decided to seed from the posterior midline cortex for three main reasons. First, unlike the associations with the frontal cortex, associations between the hippocampal system and the posterior midline cortex develop relatively early^41^. Second, adult cortical networks DN-A, DN-B and PMN/SAL are known to occupy known and anatomically adjacent regions in the posterior midline cortex, as shown consistently across different datasets. Third, we wanted to examine neocortical network representations in the hippocampal-entorhinal system using seeds that are approximately equally distant from the hippocampal-entorhinal system^42–44^. Since there is no consensus pertaining to the cortical network organization of the infant brain, each network identification was based on known network components in the adult brain spanning multiple neocortical areas as well as the hippocampus and the entorhinal cortex. After generating hypotheses for distinct cortical networks and their selectively associated distributed regions, the anatomical dissociations between candidate networks were then formally tested using independent validation datasets (other half of the data for each infant).

The whole-brain topography associated with seeds placed in the infant retrosplenial cortex (dorsal part of the posterior midline cortex - associated with the DN-A in adults) is shown on representative slices in Figure 2, upper row (see Figure S2 for other participants and Figure S3 for group-level surface representations of whole-brain connectivity patterns). In all infants, in addition to the retrosplenial cortex, the whole-brain topography included the inferior parietal lobules, dorsolateral frontal cortex and ventromedial prefrontal cortex. Importantly, in all infants, seeds placed in the retrosplenial cortex displayed connectivity with the anterior hippocampus and the most medial part of the entorhinal cortex positioned closest to the hippocampus, likely overlapping at least partially with pre/parasubiculum (Figure 3, upper row; see Figure S4 for other participants). Overall, the distributed connectivity patterns of the infant retrosplenial cortex strongly corresponded to the topography of the adult cortical network DN-A^24,27^.

**Figure 2.**
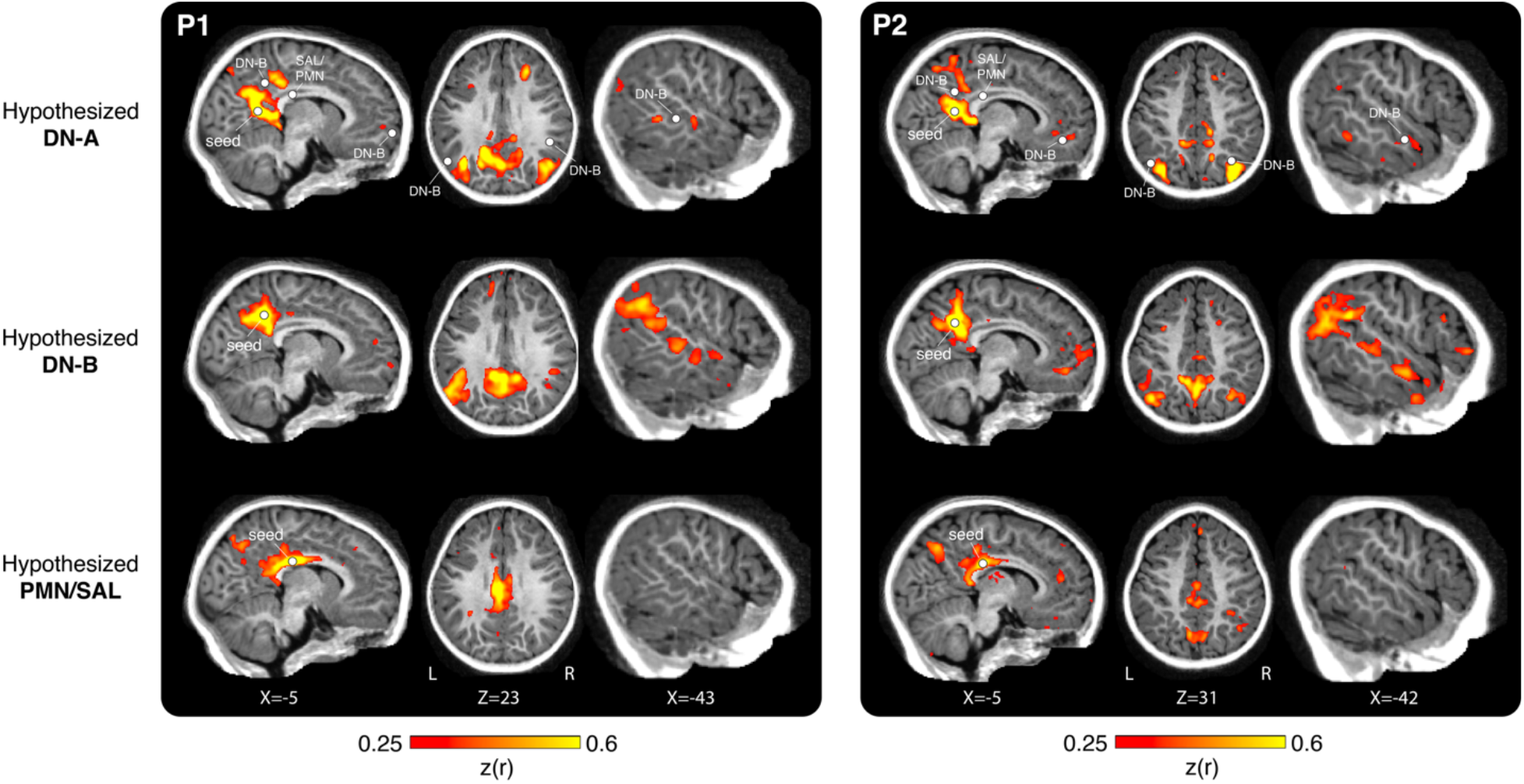
Network discovery. Using seeds placed in the posterior midline cortex we identified candidate networks DN-A (top row), DN-B (middle row) and PMN/SAL (bottom row) in the infant brain. For each participant, representative sagittal, axial and temporal slices are presented; all networks are presented on the same slice for each orientation. Note the close anatomical position of the DN-A, DN-B and PMN/SAL seeds indicated by circles in the top row. All three networks resembled their adult counterparts and occupied anatomically adjacent brain regions indicated by circles in the parietal lobe, ventral medial prefrontal cortex, and lateral surface of the temporal lobe, defined as the local maximum Fisher transformed Pearson correlation coefficient with the posterior midline seeds for each network in each presented brain slice. These and other regions were then submitted for an independent validation test, results of which are shown in Figure 4. See Figure S2 for data from other participants and Figure S3 for group-level surface representations of the whole-brain connectivity patterns.

**Figure 3.**
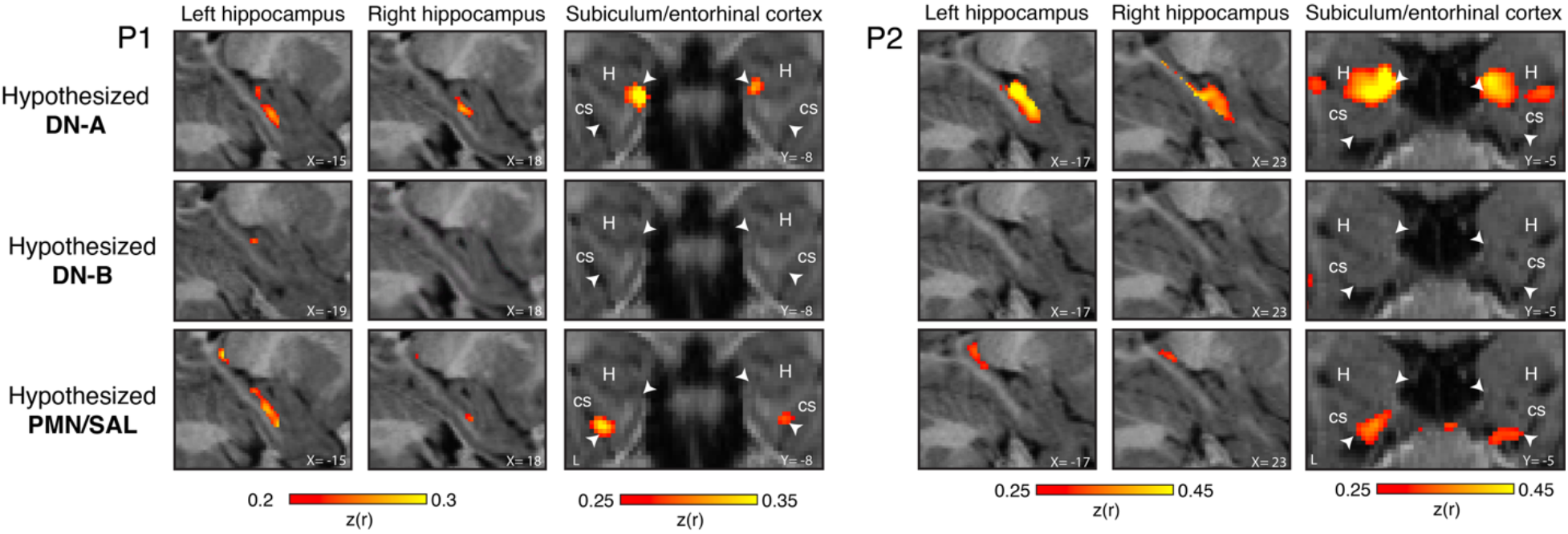
Network discovery in the MTL. Using the posterior midline seeds shown in Figure 2, we examined the associations of the candidate networks DN-A (top row), DN-B (middle row) and PMN/SAL (bottom row) with the infant MTL structures, particularly the hippocampus and the entorhinal cortex. For each participant, representative sagittal and temporal slices are presented; all networks are presented on the same slice for each orientation, except for the DN-B associations with the left hippocampus in P1. The infant cortical network DN-A was associated with the anterior hippocampus and the most medial part of the entorhinal cortex, likely overlapping at least partially with the pre/parasubiculum, similar to adults. The infant cortical network PMN/SAL was associated with the posterior hippocampus and the most lateral part of the entorhinal cortex, similar to adults (in P1 associations with anterior hippocampus were observed as well). In contrast to adults, associations of the infant DN-B with the MTL structures were minimal – weak associations with the hippocampus were observed only in P1 (depicted at the left hippocampus sagittal slice) and P5; DN-B associations with the entorhinal cortex were meager in all of the infants in our sample. The putative medial-lateral extent of the entorhinal cortex is marked with white arrowheads. See Figure S4 for data from other participants.

The whole-brain topography associated with seeds placed in the infant caudal posterior cingulate cortex (associated with the DN-B in adults) is shown on representative slices in Figure 2, middle row (see Figure S2 for other participants). These seeds resulted in whole-brain connectivity patterns that were consistent across all infants and, in addition to the posterior cingulate cortex, included the inferior parietal lobules, dorsolateral frontal cortex, ventromedial prefrontal cortex and the lateral surface of the temporal lobe. Notably, and in a striking contrast to the seeds placed only a few millimeters ventrally in the retrosplenial cortex, the seeds in the posterior cingulate cortex showed weak and no consistent connectivity with the hippocampus and the entorhinal cortex (Figure 3, middle row; see Figure S4 for other participants; see validation analysis below). Apart from the lack of consistent connectivity with the MTL observed using the discovery datasets, the overall connectivity patterns of the infants’ posterior cingulate cortex strongly corresponded to the topography of the adult cortical network DN-B^24,27^. Most of the cortical regions associated with seeds placed in the retrosplenial and posterior cingulate cortex occupied anatomically neighboring regions in an interdigitated fashion. This provided further support that the different posterior midline seeds are associated with distinct cortical networks^24,25^.

The whole-brain topography associated with seeds placed in the rostral posterior cingulate cortex (associated with the PMN/SAL in adults) is shown on representative slices in Figure 2, bottom row (see Figure S2 for other participants). In addition to the posterior cingulate cortex, the whole-brain topography included the intraparietal sulcus and ventral precuneus. Importantly, the rostral posterior cingulate cortex seeds were associated with the posterior hippocampus (in P1 also the anterior hippocampus) and the most lateral part of the entorhinal cortex positioned closest to the perirhinal cortex (Figure 3, bottom row; see Figure S4 for other participants). Even though these connectivity patterns were consistent across all infants in our study and overall resembled the adult cortical network PMN/SAL, important network components that are present in adults, especially the frontal components, were not present in infants. Specifically, the adult cortical network PMN/SAL has pronounced components in the anterior cingulate, insula and prefrontal cortex^18,27,45^, which were not present in any of the infants.

To verify that the networks we identified in the infant brain using discovery datasets and a seed-based approach correspond, at least partially, to their adult counterparts, we calculated dice coefficients between binarized infants’ discovery connectivity maps and distributed networks in DU15 parcellation^27^. This analysis showed that all networks of interest in the infant brain had the maximum dice coefficient with the corresponding network in the adult brain, except for PMN/SAL in P3 and P4 (Figure S5; for these participants, dice coefficient for PMN/SAL were second largest). Since dice coefficients for the putative SAL/PMN were much lower than for the putative DN-A and DN-B, we calculated dice coefficients separately for the anterior and posterior parts of the network (see Methods for more details). Consistent with the lack of frontal PMN/SAL components in the infants compared to the adult brain, in all participants dice coefficients were larger for the posterior PMN/SAL part, further suggesting that frontal components of this network might still be immature in infancy.

### Validation of infant distributed cortical networks and their associations with the hippocampal-entorhinal system

After generating hypotheses about distinct cortical networks in the infant brain and their differential association with the hippocampal-entorhinal system, these hypotheses were submitted to a replication test using independent validation datasets in each individual infant. For testing the dissociation between the hypothesized DN-A and DN-B, dissociation regions were defined in the parietal lobe, dorsolateral frontal cortex, ventromedial prefrontal cortex, lateral surface of the temporal lobe and parahippocampal cortex. For testing the dissociation of the hypothesized cortical network PMN/SAL, dissociation regions were defined in the precuneus and intraparietal sulcus. For testing the dissociation between the cortical networks DN-A and PMN/SAL in the hippocampal-entorhinal system, dissociation regions were defined in the posterior and anterior hippocampus, as well as in the most medial and lateral parts of the entorhinal cortex. All dissociation regions we defined using the discovery dataset in each infant. With this, we aimed to test whether the a priori posterior midline seeds used for the discovery datasets and those brain regions that maximize network separation (both in the distributed neocortex and the MTL) would show similar connectivity patterns in the validation datasets.

In the neocortex, DN-A and DN-B showed significant dissociations across all *a priori* defined brain regions, except for the lateral surface of the temporal lobe in P4 (Figure 4; tested using two-way and one-way repeated measures ANOVA; all significant p-values < 0.05). Additionally, all two-way interactions showed a cross-over effect, except for the ventromedial prefrontal cortex in P3. For the cortical network PMN/SAL, only P1 and P2 showed significant preferred connectivity of the posterior cingulate seed with both the precuneus and intraparietal sulcus (defined *a priori;* all p-values < 0.05, tested using one-way repeated measures ANOVA). Participants P3, P4, and P5 showed mixed dissociation of the cortical network PMN from DN-A and DN-B, ranging from no preferred connectivity with the posterior cingulate seed at all in the participant P5, to only intraparietal sulcus showing preferred connectivity in the other two participants (P3 & P4; Figure 4). Finally, we performed a replication analysis dissociating networks DN-A and DN-B at the neocortex using different seeds in the parietal cortex (see Methods for more details).

**Figure 4.**
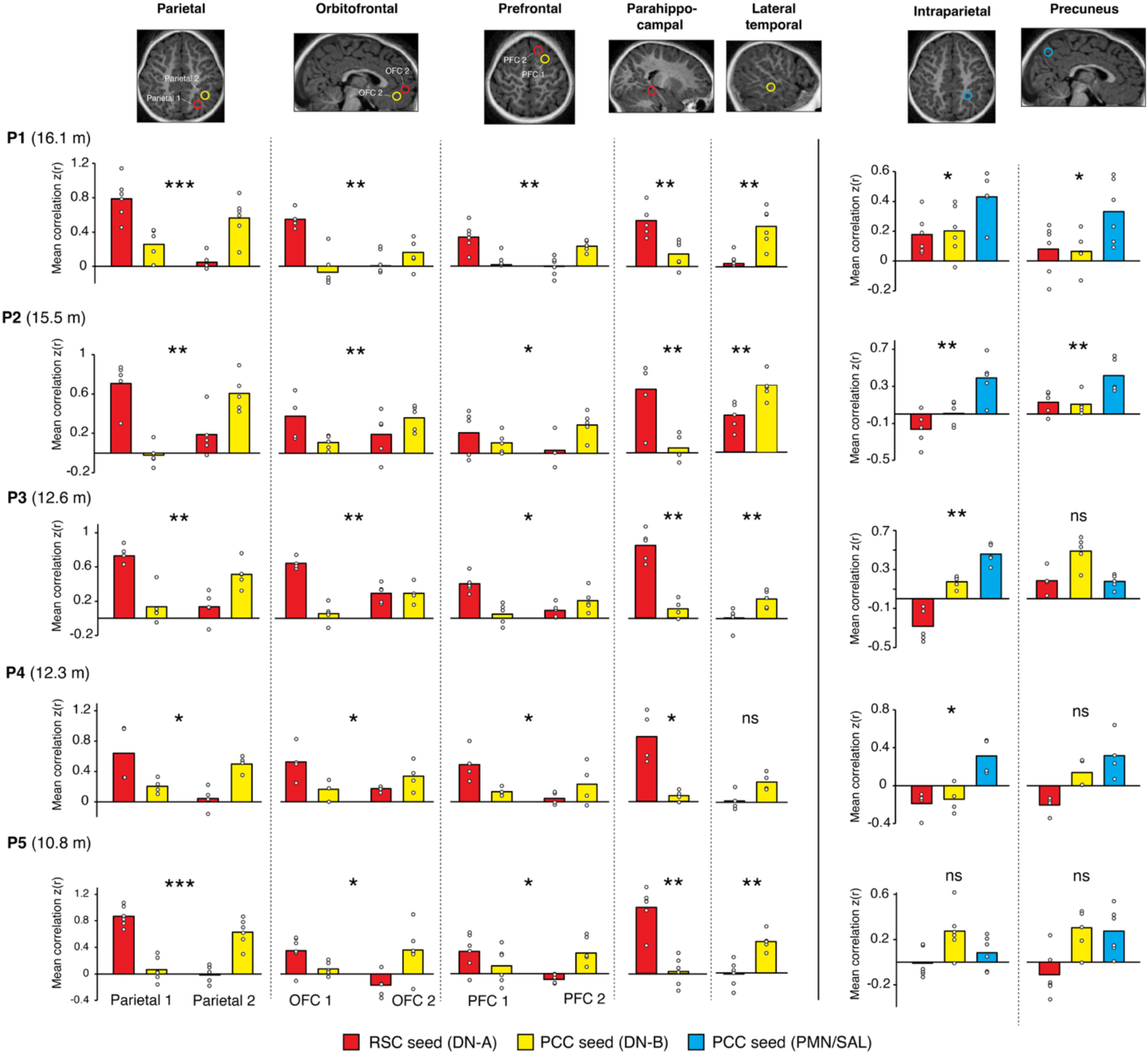
Network validation in the neocortex. Using independent validation datasets, in each infant we tested the hypotheses that distinct posterior midline seeds associate with distinct brain regions throughout the cortex, including the hippocampus and the entorhinal cortex. Importantly, all brain regions used for the validation analysis were defined *a priori* using discovery datasets. The validation analyses showed that distinct regions across the cortex correlated differently with distinct posterior midline seeds, and these distinctions were statistically significant (all significant p < 0.05), thus supporting the hypothesis of the existence of distinct cortical networks in the infant brain, corresponding to the topography of the adult DN-A, DN-B and partially PMN. Note that while cortical networks DN-A and DN-B showed robust dissociations in all infants (except for the lateral surface of the temporal lobe in P4), the cortical network PMN was fully dissociated only in the oldest infants in our sample - P1 and P2. Bars are mean correlations across all available validation dataset runs, presented as individual dots. Representative example dissociation regions are shown at the top. Numbers in parentheses represent the age of each infant, m – months. ∗p < 0.05, ∗∗p < 0.01, ∗∗∗p < 0.001.

In the entorhinal cortex, in turn, the cortical networks DN-A and PMN/SAL showed significant dissociations in all infants (Figure 5; tested using two-way repeated measures ANOVA; all p-values < 0.05). Similarly, the hippocampus also showed a significant dissociation between DN-A and PMN/SAL along its longitudinal axis in all infants except for participant P5 (tested using two-way repeated measures ANOVA; all significant p-values < 0.05). All significant two-way interactions between the DN-A and PMN/SAL regions in the hippocampus and the entorhinal cortex showed a cross-over effect (Figure 5).

**Figure 5.**
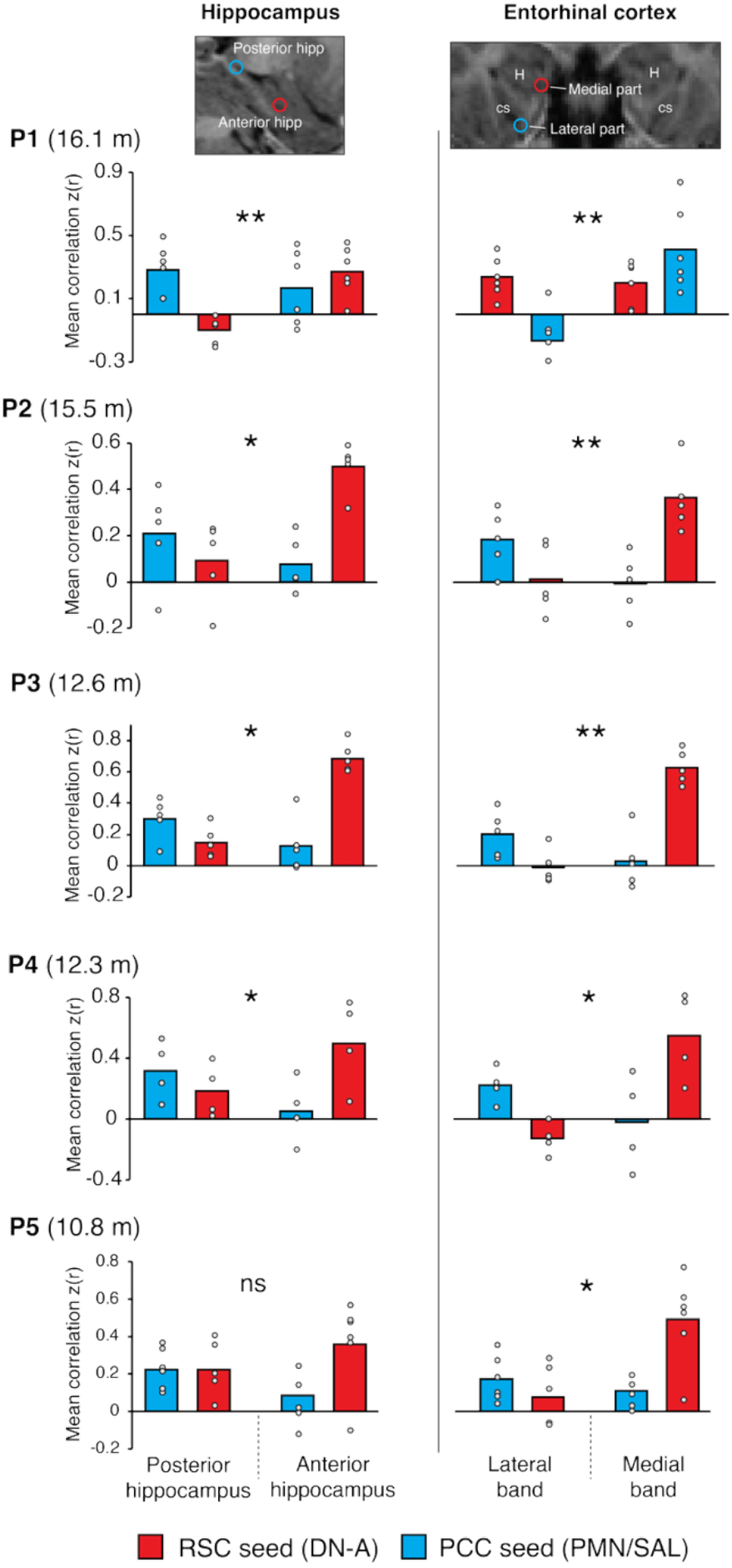
Network validation in the medial temporal lobe. Using independent validation datasets, in each infant we tested the hypotheses that distinct posterior midline seeds associate with distinct parts of the hippocampal long axis and distinct parts of the entorhinal cortex. All brain regions used for the validation analysis were defined *a priori* using discovery datasets. The distinct posterior midline seeds, representing hypothesized cortical networks DN-A and PMN, were associated with different parts of the infant hippocampal-entorhinal system (all significant p < 0.05). These results support the hypothesis that the most medial part of the entorhinal cortex is associated with the cortical network DN-A and the most lateral part of the entorhinal cortex is associated with the cortical network PMN/SAL already by approximately one year of age. Additionally, we find that in all infants except for the youngest, P3, the hippocampal long axis was functionally dissociated: the anterior hippocampus was preferably associated with the DN-A and the posterior hippocampus with the PMN. Bars represent mean correlations across all available validation dataset runs; dots represent individual validation datasets. Numbers in parentheses represent the age of each infant, m – months. Representative example dissociation regions are shown at the top. ∗p < 0.05, ∗∗p < 0.01.

To validate the weaker associations between the cortical network DN-B and the infant hippocampal-entorhinal system that we observed using discovery datasets, we calculated the percentage of hippocampal and entorhinal cortex voxels that passed the noise threshold level (z(r)>0.15) for each network of interest using validation datasets. Importantly, in addition to the original posterior midline seeds, in each infant, we calculated the percentage of threshold-passing voxels using seeds placed in distributed dissociation regions spanning inferior parietal cortex, orbitofrontal cortex, and dorsolateral prefrontal cortex (all defined *a priori* using discovery datasets; all seeds were defined by having maximum local connectivity with the original posterior midline seeds). As shown in Figure 6, across participants, the average percentage of threshold-passing voxels associated with DN-B was 6.4% in the hippocampus and 2.3% in the entorhinal cortex. For DN-A, the average percentage was significantly higher – 41.8% in the hippocampus and 23.9% in the entorhinal cortex (paired t-test, hippocampus - t_(4)_=15.31, p<0.001, entorhinal cortex - t_(4)_=36.03, p<0.001). For reference, the average percentage of threshold-passing voxels associated with PMN/SAL was 24.8% and 22.4% in the hippocampus and the entorhinal cortex, respectively, significantly higher compared with the DN-B (hippocampus - t_(4)_=6.23, p<0.01, entorhinal cortex - t_(4)_=6.67, p<0.01; see Figure S6 for the same analysis using different noise-level thresholds).

**Figure 6.**
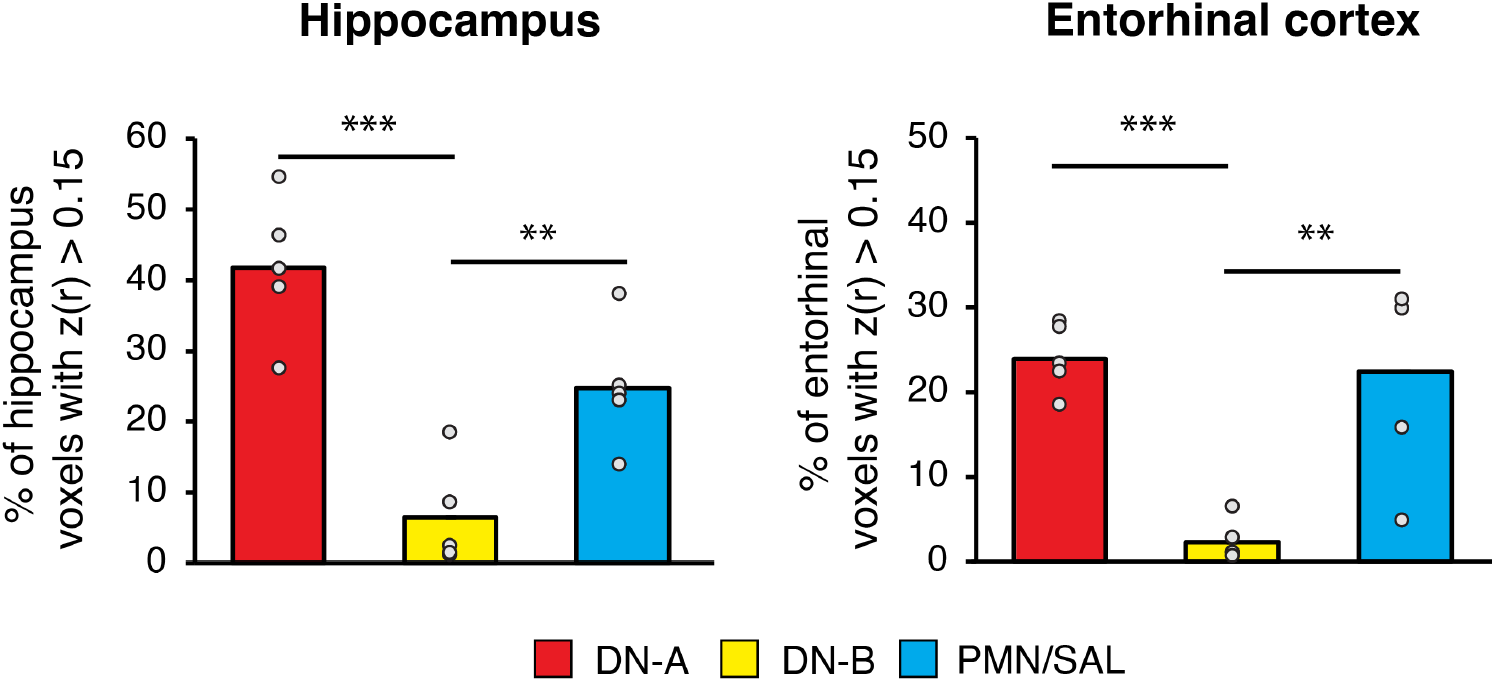
Validation of weaker DN-B associations with the infant MTL. Using independent validation datasets, we examined the average percentage of threshold-passing voxels in the hippocampus and the entorhinal cortex, associated with DN-A, DN-B and PMN/SAL. All brain regions used for this validation analysis were defined *a priori* using discovery datasets. Associations with the cortical network DN-B were significantly lower than with DN-A and PMN/SAL (all p-values < 0.01). Bars represent mean percentage of threshold-passing voxels across all seeds, averaged across participants, presented as individual dots. ∗∗p < 0.01, ∗∗∗p < 0.001. See Figure S6 for the same analysis using different noise-level thresholds.

In addition to formal statistical testing, we performed a “back projection” analysis using the validation datasets. We placed seeds (defined *a priori* using the discovery datasets) in the most medial and lateral parts of the entorhinal cortex that showed maximum associations with the cortical networks DN-A and PMN/SAL in the discovery datasets, respectively, and calculated whole-brain “model-free” connectivity maps. The aim of this analysis was to test whether “back projection” topography would align with the expected parts of the hippocampal long-axis, as well as with DN-A and PMN/SAL topographies. As shown in Figure 7 (see Figure S7 for data from other participants), seeds placed in the most medial part of the entorhinal cortex (potentially overlapping at least partially with the pre/parasubiculum) resulted in whole-brain topography, which aligned with the cortical network DN-A, including the anterior hippocampus. Seeds placed in the most lateral part of the entorhinal cortex resulted in very different whole-brain topography resembling the cortical network PMN/SAL, especially the posterior cingulate component, including the hippocampus, extending to its most posterior parts. These results provide further support that distinct parts of the infant entorhinal cortex are associated with distinct parts of the hippocampal long axis and different neocortical networks.

**Figure 7.**
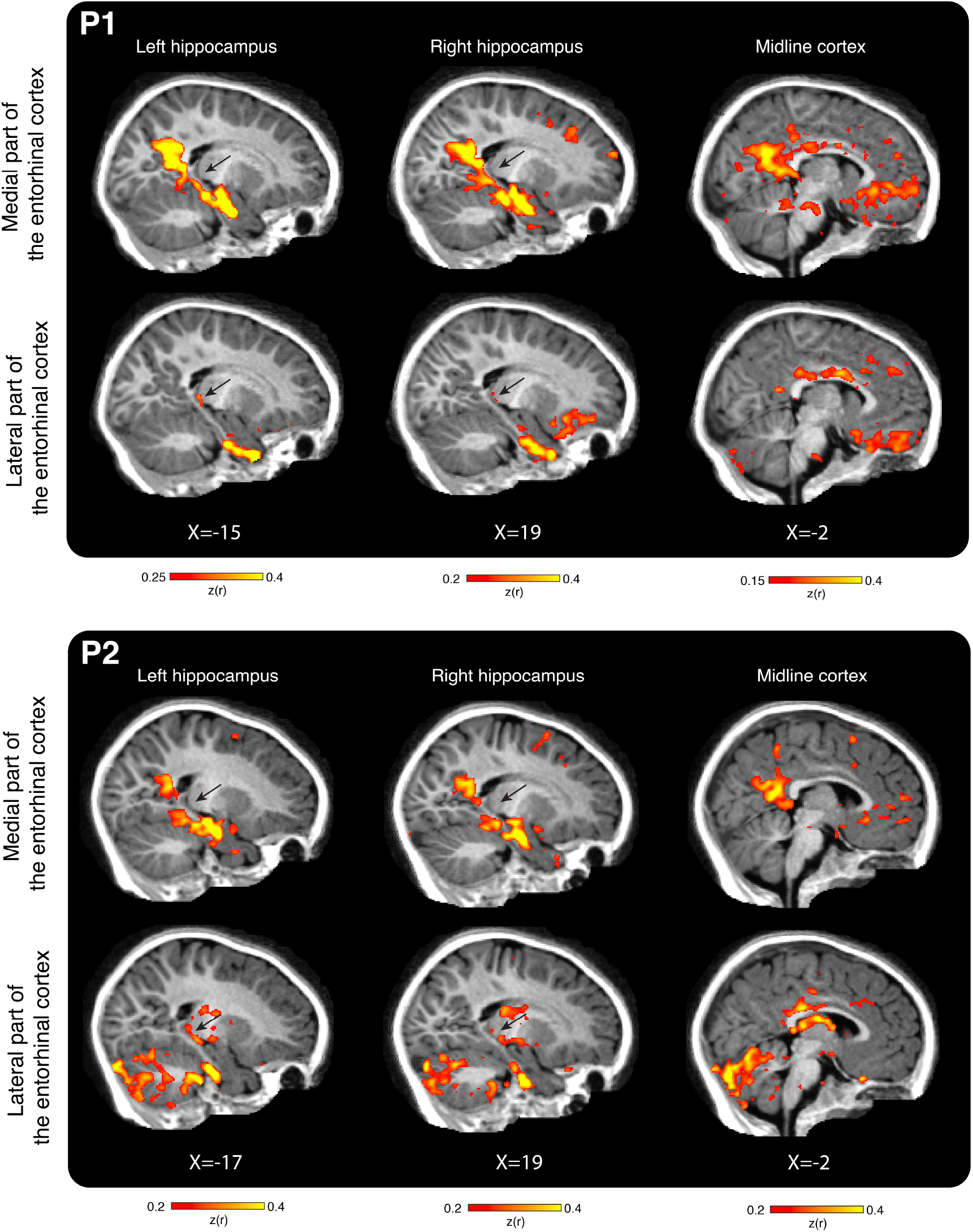
“Back projections” from the entorhinal cortex. In addition to formal statistical testing, we examined whether seeds placed in distinct parts of the infant entorhinal cortex, informed by the discovery dataset, would associate with distinct parts of the hippocampal long axis and distinct cortical networks in the validation dataset. We found that seeds placed in the most medial and lateral parts of the infant entorhinal cortex revealed whole brain topographies corresponding to infants’ DN-A and partially to PMN/SAL, including anterior and posterior hippocampus (marked with black arrows) respectively. These results provide further support for the existence of distinct entorhinal-hippocampal circuits in the infant brain that associate distinct parts of the entorhinal cortex with distinct parts of the hippocampus. See Figure S7 for data from other participants.

The finding that the cortical network DN-B showed significantly weaker connectivity with the MTL in our sample of infants aged 10 to 16 months raises the question of how this network matures beyond 16 months as important cognitive milestones in social cognition and episodic memory development are reached in the second year of life. To investigate this, we reinvited the infants after approximately 9 months following the original testing session to explore the longitudinal development of MTL connectivity with DN-A, DN-B and PMN/SAL within individuals.

### Exploratory analysis: development of DN-B representations in the hippocampal-entorhinal system

Based on our finding of weak MTL connectivity with the DN-B by 16 months of age, we next examined the potential developmental trajectory of the hippocampal-entorhinal system across the second year of life by collecting exploratory longitudinal precision fMRI data from the same infants approximately 9 months after the initial session. Longitudinal precision imaging in infants poses substantial practical challenges, owing to the large amounts of high-quality data required at each time point, leading to extraordinarily high attrition-rates, as well as reduced deep daytime sleep across the second year of life. We obtained sufficient, high-quality fMRI data from two of the participants (P1 and P2; Table 1), who were approximately two years of age at the second time point (t2, see Methods for details). To ensure comparability across time, we used the same DN-A, DN-B and PMN/SAL posterior midline seed regions defined using the discovery datasets from time point one (t1). This longitudinal individualized precision dataset enabled us to perform an exploratory analysis to assess how associations of the cortical networks DN-A, DN-B and PMN/SAL with the hippocampal-entorhinal system evolve in the second year of life.

First, we found that for the cortical network DN-B, the percentage of hippocampal voxels that passed the noise threshold of z(r) > 0.15 significantly increased between time points t1 and t2 from 4.4% to 25.9% in P1 and from 3.6% to 25.7% in P2 (Figure 8A, top row, marked with white arrows; difference of two proportions test, p-values < 0.001). Importantly, using the same seed in the posterior cingulate cortex as at time point t1, we found that at time point t2 the cortical network DN-B in both infants was associated with the part of the entorhinal cortex located between the medial and lateral parts associated with DN-A and PMN/SAL respectively, aligning with the adult topography (Figure 8D, marked with white arrows). To compare the association of the DN-B with the entorhinal cortex between time points t1 and t2, we examined the percentage of the entorhinal voxels associated with the DN-B that passed the noise threshold at each time point. We found that the percentage of the entorhinal voxels associated with the DN-B significantly increased between t1 and t2 from 0% to 4.7% in P1 and from 0.15% to 13.94% in P2 (Figure 8D, p-values < 0.001).

**Figure 8.**
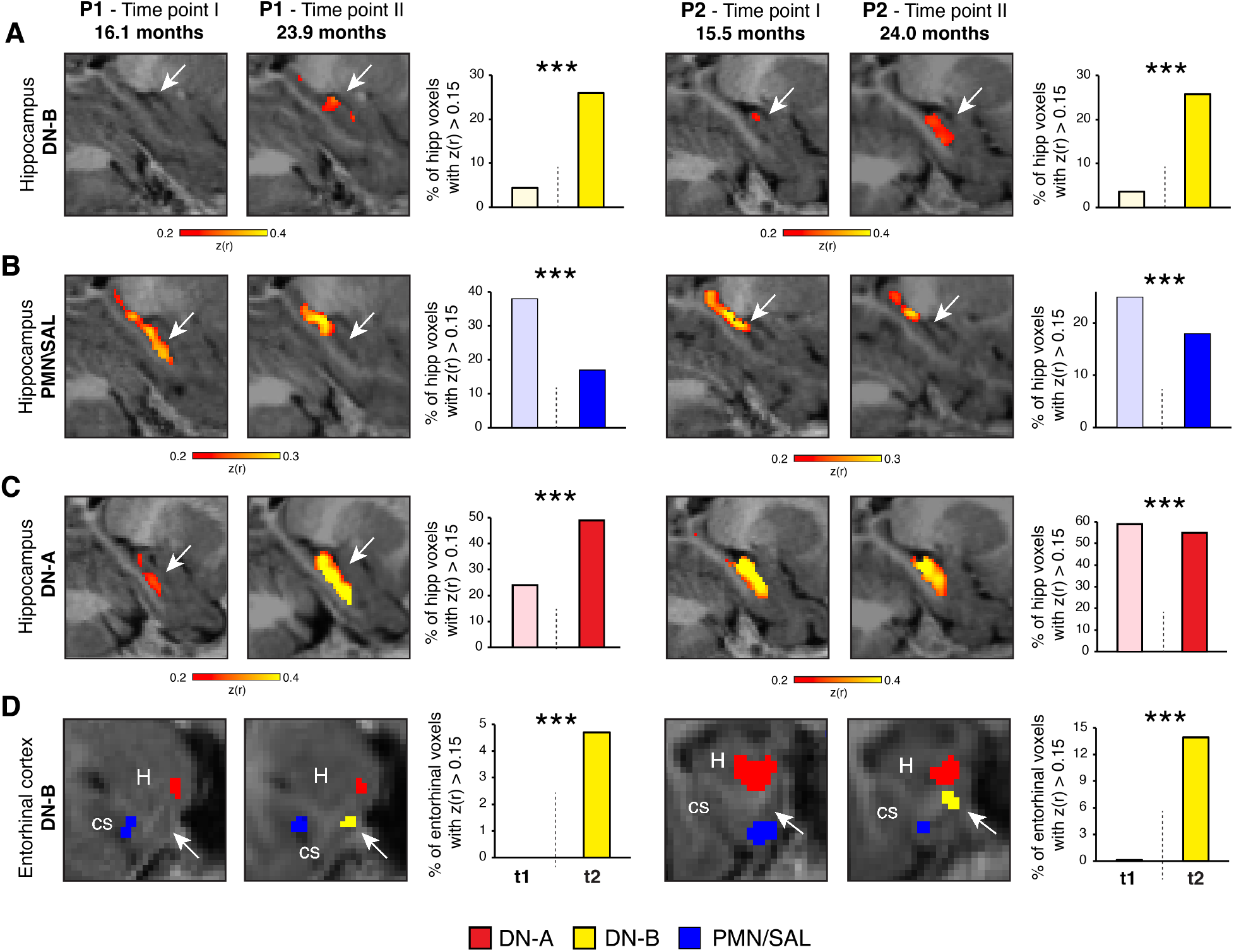
Exploratory analysis examining longitudinal changes in cortical network associations with the infant MTL. We leveraged longitudinal precision data that we collected from two infants to examine how associations of the cortical networks DN-A, DN-B and PMN/SAL with the hippocampus and the entorhinal cortex change between 15 and 24 months of age. **A**. First, we observed that the percentage of hippocampal voxels that pass the noise correlation threshold increased from approximately 4% to 27% for both P1 and P2. **B.** Second, we observed that the percentage of hippocampal voxels associated with the cortical network PMN that pass the noise correlation threshold decreased from 38% to 17% in P1 and from 25% to 18% in P2 (marked with white arrows, p < 0.001). Importantly, this decrease resulted in more posteriorly localized hippocampal associations with the PMN/SAL, similar to that of adults. **C.** Third, the change of the percentage of hippocampal voxels associated with the cortical network DN-A that pass the noise correlation threshold was not consistent across P1 and P2. While in P1, the percentage of hippocampal voxels increased, in P2 it decreased (albeit modestly; marked with white arrows, both p < 0.001). **D**. Finally, our results indicate a significant increase in the percentage of entorhinal cortex voxels that pass the noise correlation threshold from 0% to 4.7% in P1 and 0.015% to 13.94% (marked with white arrows, both p < 0.001; note that associations of each cortical network with the entorhinal cortex were thresholded to display only the local maximum correlation coefficients; for clarity, only entorhinal voxels are shown). Se Figure S8 for the same analyses using different noise thresholds.

For the PMN/SAL, we found that the percentage of hippocampal voxels associated with the PMN/SAL seed in posterior cingulate cortex that passed the noise threshold of z(r) > 0.15 decreased between time points t1 and t2 from 38% to 17% in P1 and from 25% to 18% in P2 (Figure 8B, middle row, marked with white arrows, p-values < 0.001). Importantly, this decrease reflected an increasingly selective association of the PMN/SAL with voxels in the more posterior parts of the hippocampus, corresponding to a more adult-like organization^15,19,20^. These results indicate that, in the second year of life, hippocampal connectivity of the PMN/SAL becomes more topographically restricted.

Changes in hippocampal associations with the cortical network DN-A between t1 and t2 were not consistent across infants. While in P1 we observed an increase in the percentage of hippocampal voxels associated with DN-A that passed the noise threshold of z(r) > 0.15 (24% in t1 to 49% in t2; p < 0.001), in P2 we observed a modest decrease in the percentage of hippocampal voxels that passed the noise threshold of z(r) > 0.15 (59% in t1 to 55% in t2; p < 0.001; Figure 8C, bottom row, marked with white arrows; see Figure S8 for the same analyses using different noise thresholds and Figure S9 for the distributed topographies of DN-B, PMN/SAL and DN-A at time point 2).

Taken together, our exploratory longitudinal findings suggest that the connectivity of the DN-B with the human hippocampal-entorhinal system may increase in the second year of life. The PMN/SAL, in turn, shows more posteriorly localized associations in the hippocampus, corresponding to a more adult-like connectivity pattern. Together, these findings potentially point to a gradual integration and refinement of distinct cortical networks’ associations with the human hippocampal-entorhinal system during the second postnatal year.

## Discussion

In the current study we collected precision fMRI data from typically developing infants between 10 and 24 months of age to study the development of the hippocampal-entorhinal system and its integration with distributed cortical networks. Our results point to three organizational principles of the developing hippocampal-entorhinal circuitry in humans. First, already by 10 months of age, the infant cortical network DN-A is organized similar to the adult DN-A, including associations with the anterior hippocampus and the most medial part of the entorhinal cortex. Second, while the DN-B is already fully differentiated from the DN-A in the neocortex, its associations with the hippocampal-entorhinal system are significantly weaker by 16 months of age compared with DN-B and our exploratory findings in two infants suggest that these associations may significantly increase towards the end of the second postnatal year. Third, the hippocampal long axis and the entorhinal medial-lateral axis are already differentiated by 1 year of age, showing preferred associations of the anterior and posterior hippocampus and the most medial and lateral part of the entorhinal cortex with the cortical networks DN-A and PMN/SAL, respectively. Our exploratory findings suggest that the association of the cortical network PMN/SAL with the hippocampus may become more topographically restricted to the posterior hippocampus in the second year of life, corresponding to the adult-like topography, suggesting that hippocampal long-axis differentiation continues well after 1 year of age.

### Relations to prior studies

Previous precision fMRI mapping studies in human adults provided critical insights into fine-grained organizational principles of distributed cortical networks^24–27,46,47^, hippocampus^16,19,20^, entorhinal cortex^15,16^, and various subcortical structures, such as the amygdala^17,48^ and cerebellum^49,50^. Historically, it was challenging to facilitate these discoveries using traditional group-average approaches where fine and idiosyncratic anatomical differences are blurred across individuals. The challenge of group averaging is particularly pronounced in developmental studies, where infants’ brain size is much smaller than adults^51^ and data quality is notoriously low^36,52^. Therefore, within individual, precision mapping of the infant brain is a powerful means for studying the organizational principles of the human brain throughout cognitive development^53–57^. Indeed, our precision fMRI findings in infants dissociate distinct developmental profiles of different parts of the infant hippocampal-entorhinal system and point to a gradual development of the human hippocampal-entorhinal-cortical circuitry.

A previous study in human adults showed that similar to all studied mammalian species, the human entorhinal cortex is organized into band-like connectionally defined zones^16^ that divide the entorhinal cortex along its rostro-caudal axis into three medial-lateral bands – the medial band (closest to the hippocampus; associated with the anterior hippocampus and DN-A), the lateral band (positioned near the collateral sulcus; associated with the posterior hippocampus and PMN/SAL), and the intermediate band (positioned in between the other two band; associated with the anterior hippocampus and DN-B). Importantly, these entorhinal bands associate with different parts of the hippocampal long-axis – the medial band is preferably associated with the anterior hippocampus (ventral in rodents), the intermediate band is associated with the intermediate hippocampus, and the lateral is associated with the posterior hippocampus (dorsal in rodents).

Based on our infant fMRI data, we suggest that the most medial and lateral parts of the entorhinal cortex correspond at least partially to the medial and lateral entorhinal bands, respectively. This association is supported by the relative anatomical location along the entorhinal medial-lateral axis, as well as differential associations with the hippocampal long-axis and with distinct cortical networks – DN-A and PMN/SAL. Associations of the DN-B with the hippocampus and the entorhinal cortex are significantly weaker around one year of age, and they were apparent only in the data collected in the later half of the second postnatal year. These DN-B associations were localized to the parts of the entorhinal cortex positioned between these associated with DN-A and PMN/SAL, suggesting that this may correspond to the intermediate entorhinal band, which is associated with the DN-B in adults^16^.

Previous infant studies showed that the canonical default network is already present shortly after birth^43,44,58–60^. Based on the undifferentiated monkey homologue of the canonical human default network^21,61^, an intriguing developmental hypothesis suggested that a prototypical, undifferentiated default network exists in earliest developmental stages that undergoes experience-dependent differentiation into its subnetworks later in human life^21^. It was suggested that their differentiation may be facilitated by differential associations with the MTL structures, which may provide anatomically selective input to parts of an initially undifferentiated default network. Our results indicate that this differentiation is already established by the end of the first postnatal year, thus, motivating further research into the detailed organizational properties of the neonatal default network at even earlier ages^55^.

The later developing association between the infant MTL and DN-B was in contrast to the development of the association between the MTL and PMN/SAL. While the neocortical differentiation of the PMN/SAL still seemed immature in the studied age range, it was already associated with the hippocampus and entorhinal cortex by around 1 year of age. In line with findings in adults, PMN/SAL showed preferred associations with the posterior hippocampus and the lateral entorhinal band. At the neocortical level, infants’ PMN/SAL was missing several components that exist in adults, such as the anterior cingulate, prefrontal and insular components^18,27,45^. While the incomplete architecture of PMN/SAL in infancy is consistent with previous studies^43,54,60,62,63^, it may suggest that this network should not be treated as a full homologue of the adult PMN/SAL. Indeed, some distributed cortical networks develop remarkably late during infancy, and while the canonical default network is already present in its adult form by 12 months of age, other networks at that age lack either posterior or anterior components^58,64^. In our data, long-range connections were absent specifically for the PMN/SAL and not for the DN-A and DN-B, suggesting that the lacking frontal components of PMN/SAL cannot be solely attributed to generally weaker long-range connections in the infant brain.

It is important to mention that following pioneering group-level studies into the human hippocampal region in adults, the human MTL is traditionally associated with two memory systems – the anterior memory system, associated with the perirhinal cortex, and the posterior memory system, associated with the parahippocampal cortex^65–67^. Previous inquiries into the functional neuroanatomy of the human entorhinal cortex further associated different subregions of the entorhinal cortex with the perirhinal and parahippocampal cortex^68^. Recent breakthroughs into the functional neuroanatomy of the anterior medial temporal lobe using precision neuroimaging at ultra-high field 7-Tesla fMRI revealed the remarkably complex organization of the human entorhinal cortex and its whole brain distributed circuitry^15–17^. Therefore, we suggest that what is traditionally considered as the posterior and anterior memory systems, might overlap at least partially with the canonical default network, which in turn, could be further divided at least into DN-A, DN-B, and often PMN/SAL^69^ (see also Ref^70^ for an attempt to harmonize network naming conventions).

### Relations to rodent and monkey development

Studies in neonate rodents and monkeys show that adult-like patterns of entorhinal-hippocampal connectivity are already present shortly after birth^71,72^ and can even be observed at very early stages of fetal development. Nevertheless, different components of the hippocampal-entorhinal system mature at different postnatal stages. For example, in rodents, the medial entorhinal cortex matures earlier than the lateral entorhinal cortex^2^ and the dorsal/caudal part of the medial entorhinal cortex matures earlier than its ventral/rostral part^73^. Similar findings were reported in monkeys, showing that posterior subdivisions of the monkey entorhinal cortex mature earlier than more anteriorly positioned subdivisions^13^. In line with the modular development of the animal entorhinal cortex, hippocampus as well shows differential developmental trajectories along multiple anatomical axes, including the longitudinal axis^74,75^. Taken together, these animal findings support the notion that different hippocampal-entorhinal-cortical circuits show different developmental profiles.

In rodents, the band-like organization of the entorhinal cortex develops very early with all three entorhinal bands being integrated into the entorhinal-hippocampal circuitry long before eyes and ear canal opening^76^. While our knowledge pertaining to the development of neocortical connections of different entorhinal bands in animals is scarce, our results suggest that neocortical and hippocampal associations with the intermediate entorhinal band (associated with the DN-B) may emerge latest in human development. Furthermore, connectivity of the entorhinal cortex with the posterior midline cortex is remarkably similar between adult monkeys and 1-year-old infant humans^77–79^. This correspondence between ontogeny and phylogeny suggests that the MTL circuitry associating at least parts of the cortical network DN-B with the hippocampal-entorhinal system may be evolutionary novel in humans^15^, potentially underpinning unique properties of human hippocampus-dependent cognition^80^.

### Functional implications

Previous studies in adults have shown that the cortical networks DN-A, DN-B and PMN support different cognitive functions. The DN-A and DN-B are implicated in internally-oriented processing, in particular episodic simulation for the DN-A and social cognition and mentalizing for the DN-B^26^. The PMN/SAL, in turn, is implicated in tasks involving processing external information^27^, such as novelty detection^20^ and recognition memory^32^. Our current findings indicate that infants’ hippocampal-entorhinal system is associated with DN-A and PMN/SAL already by one year of age. The adult-like topography of the DN-A including its connectivity with the MTL may provide the anatomical scaffold for hippocampal encoding of memories recently observed from around 1 year of age^11^. Further, our findings align with relational binding^81–83^ and spatial capacities^84^ present in infants around that age, which are considered precursors of mature episodic memory and navigation^85^, supported by the DN-A in adults^86^.

Potential functional insights emerge from our exploratory analysis examining the longitudinal trajectory of PMN/SAL and particularly DN-B associations with the infant hippocampal-entorhinal system. In animals, the gradual maturation of distinct components of the hippocampal-entorhinal system is typically associated with an increase in complexity of spatial and mnemonic behaviors^73^. In humans, major breakthroughs in episodic memory and social cognition happen in the second year of life^5,6,87^. While the extent to which infants possess episodic memory remains debated, early capacities relevant to episodic memory have been demonstrated during the first 1.5 years of life, including some indication of relational memory for face-scene associations^88^ and memory for previously demonstrated action sequences^89^. More robust indicators of context-specific episodic-like memory have been reported between 1.5 to 2 years of age, including explicit recall of context-specific where-what associations^90,91^ (see Ref^6,85^ for a review).

At the same time, major advances in social cognition are achieved in the second year of life^92^. While mentalizing abilities (involving the DN-B in adults) are debated^93,94^, major developments in related precursor abilities such as joint attention^95^ and self-other distinction^92,96,97^ take place in the second year of life. In particular, between 16 and 24 months of age, there is a major breakthrough in infants’ social cognitive abilities reflected in the emergence of a self-concept^87,97,98^, which has been linked to a shift in infant memory development^92^. Specifically, in this period, linked to the development of self-concept, infants’ memory undergoes a shift towards using the self as an anchor for encoding new information^92^. Our exploratory analysis, suggests that the emerging association of the cortical network DN-B with the infant hippocampal-entorhinal system might underpin this development and may plausibly serve as a precondition for linking memories to the self, thus generating a temporal autobiographical continuity, paving the way for the developments observed in episodic-like memory towards the end of the second year of life^90,91,99^. In sum, our findings raise exciting questions for future research concerning how the developing integration of the DN-B into the hippocampal-entorhinal systems in the second year of life map onto concurrent developments in social cognition and episodic-like memory.

### Limitations

It is important to consider four main limitations of the current study. The first limitation is that we examined infants’ brain activity during natural sleep. In adults, fMRI connectivity is modulated during sleep and during different sleep stages^100–103^. Sleep is also known to modulate fMRI connectivity in infants, albeit to a weaker extent^104^; however, the effects of different sleep stages on network architecture are currently unknown in infants, inviting novel research into this important question^105^. The second limitation is that in the current study we examined only network architecture during spontaneous brain activity without imposing any task demands. It is important to remember that existence of a circuitry does not imply that the circuitry is functional^72,106^. For example, some distributed network connections are observable already during the prenatal period^107^. Furthermore, it is important to consider not only the presence of a circuitry but also interactions between different circuits as one of the driving forces underpinning the development of cognitive functions^108^. The third limitation pertains to changes in brain vasculature during the first postnatal year. Cerebral blood flow is known to increase during the postnatal period^109^ and to be associated with increased long-range fMRI connectivity^110^. However, since we observed an increase in MTL-connectivity with the DN-B and a decrease with the PMN/SAL (both reflecting more adult-like connectivity patterns), it is unlikely that the longitudinal results could be explained by developmental changes in cerebral blood flow. The fourth limitation pertains to the small number of participants that is common for precision imaging studies as these focus on dense within-individual sampling rather than on group-level inference. The amount and quality of within-individual data results in robust statistical power at the individual-infant level as well as in fine-grained insights into idiosyncratic anatomical properties and their developmental trajectory within the individual that cannot be resolved in such detail using group-averaged data.

## Conclusions

Using precision neuroimaging, we characterized distinct distributed cortical networks and their associations with the hippocampal–entorhinal system between 10 and 16 months of human life. We demonstrate that, similar to other species, the hippocampal–entorhinal system in humans develops in a modular fashion with distinct parts of the hippocampal–entorhinal circuitry developing at distinct developmental stages. In particular, our findings suggest that the association of the hippocampus and the entorhinal cortex, particularly the intermediate entorhinal band, with the cortical network DN-B—associated with social cognition—develop relatively late, pointing to a potentially distinct feature of the human hippocampal-entorhinal circuitry. We propose that this circuit may provide a developmental scaffold linking social cognition to the emergence of human episodic memory.

## Acknowledgments

The authors thank Sophie Reichard for helpful input to the manuscript. D.R., C.C.K., F.S. and C.G.W. acknowledge the support of the Max Planck Society. D.R is supported by an ERC Starting Grant (ENTONET, 101220569) and the Kavli Foundation; S.F. is supported by a Human Frontier Science Program (HFSP) Long-Term Fellowship and by the Israeli Council for Higher Education through the Bater Fellowship for Outstanding Women Doctoral Students; M.P.W. Is supported by the Kavli Foundation; C.G.W. is supported by an ERC Starting Grant (REPRESENT, 101117806).

## Data availability statement

Sharing the MRI data is not possible at this moment for data protection reasons as the data cannot be fully anonymized due to the longitudinal nature of the main study from which the infants were recruited.

## Methods

### Human infant participants

In this study we collected spontaneous fMRI data from five typically developing infants (four females, one male; age range – 10 months to 16 months). The MRI data were collected during natural sleep across multiple sessions spanning several days with the goal of collecting approximately 1 hour of spontaneous fMRI data for each infant. For three infants, data were collected across two sessions; for 2 infants data were collected across three sessions. One of the main challenges in infant MRI-based experiments is extremely low data quality due to infants’ head movement, either during natural sleep or awake states. Therefore, infants in this study were recruited from an existing large-scale group-level study, which included only one MRI session. We were particularly interested in infants who remained asleep during most of the scanning session of the original study. Families of these infants were then invited to additional sessions until approximately 1 hour of spontaneous fMRI data were collected for each infant. Recruitment for this study was done on a rolling basis and concluded once we recruited 5 infants. Importantly, additional fMRI data were collected for some of these infants after approximately eight months (see below). No power analysis was performed for determining the sample size for this study and the sample size was based on previous precision mapping studies in adults estimating functional connectivity within individuals. In total, 78 infants were screened for this study. All parents provided written consent in accordance with guidelines approved by the ethics committee of Leipzig University.

### MRI data acquisition

MR data were collected on a 3T Siemens Prisma scanner (Siemens Healthineers, Erlangen, Germany). A 20-channel commercial coil (Nova Medical, Wilmington MA, USA) was used for radio frequency transmission and reception. Each of the infant scanning sessions consisted of 4 functional runs, each lasting 12 mins, for estimating intrinsic functional connectivity. Functional imaging data were acquired using a 2D, multi-band gradient-echo echo-planar imaging (EPI) pulse sequence with the following parameters: repetition time (TR) = 2000 ms; TE = 24.2 ms; flip angle 80°; isotropic spatial resolution of 2 mm; 63 interleaved slices covering the entire cerebral cortex and the cerebellum without gaps; multiband factor of 2. The EPI sequence was a custom sequence provided by the Center for Magnetic Resonance Research (CMRR) at University of Minnesota^111^.

To estimate the distortions caused by static magnetic field (B0) inhomogeneities, the phase-encoding polarity technique was applied which combines two spin-echo EPI scans with different phase encoding directions (anterior-posterior and posterior-anterior). The field map was estimated from the difference in distortion between the two acquisitions. Field of view, imaging matrix, slice thickness and bandwidth for the Spin-Echo EPI scans were identical to the fMRI acquisition parameters. For the spin-echo scans, we used a TR of 8000 ms and a TE of 50 ms. After the initial Spin Echo EPI scan, additional scans were acquired in case infants displaced their heads during sleep to account for run specific distortions. In the beginning of the scanning session, a T1-weighted structural image was acquired using an MP2RAGE three-dimensional sequence with TR = 5000 ms; TE = 2.42 ms; spatial resolution of 1mm isotropic.

Prior to data collection, infants were put to sleep inside the scanner bore by their primary caregiver. Before falling asleep, infants were equipped with age-appropriate silicon inner-ear earplugs and if tolerated by the infant, with MRI compatible earmuffs for enhanced sound protection during the scan. After we validated that the infants fell asleep using the arm-drop-test, they were prepared for scanning by positioning their head in the scanning coil and stabilizing the infant’s position in the scanner bore with an inflatable vacuum mat to reduce motion and increase comfort. If the infant tolerated MRI compatible earmuffs prior to falling asleep, their placement was checked again before the beginning of the scan. If the infant did not tolerate the earmuffs, they were placed on the infant after falling asleep. In both cases, the infant’s head was padded with foam in the head coil. The caregiver and a member of the research team were present in the scanner room at all times during the scan. Since the data were collected during natural sleep, the goal of the research team member was to constantly monitor the infants from inside the scanner by direct visual contact. For each infant the scanning protocol began with acquiring a T1-weighted structural image, followed by acquiring an initial field-map scan. Functional data were then collected until the end of the session or until infants woke up. To account for random changes in head position during sleep, additional field-map scans were acquired in case infants spontaneously displaced their head during sleep without waking up. When infants showed a sign of waking up, the scanning procedure was immediately stopped.

### Data processing

Due to the within-individual nature of the current study, the processing of the infant fMRI data followed previous individualized preprocessing approaches^15,25,26^ and was optimized for individual-subject analysis to preserve anatomical detail across different scanning runs and sessions. In short, we calculated five spatial transformation matrices for each available BOLD run that were then combined into a single interpolation transformation applied to the original volumetric data. Matrix 1 resulted from motion correction performed by aligning all volumes within a run to its middle volume. Matrix 2 resulted from field map correction, in which every middle volume from all available BOLD runs was field map unwarped using the session-specific and run-specific field map. We used not only session-specific, but also run-specific (i.e., within sessions) field map, if the infant would displace their head without waking up. Field maps were prepared using FSL topup tool. Matrix 3 resulted from the alignment of all available BOLD runs to the mean BOLD template using a linear registration with 12 DOF. Matrix 4 resulted from the alignment of the mean BOLD template to the native T1 space. Matrix 5 resulted from the alignment of the T1 space to age-matched MNI space^112^ using a nonlinear registration. Matrices 1–5 were combined into a single interpolation transformation that was then applied to the original raw volumes from all available BOLD runs. We used this single interpolation (as opposed to serial interpolations of the data applied step-after-step) to reduce interpolation artifacts and to maintain as much as possible anatomical detail of within-participant datasets.

To prepare the data for functional connectivity analysis, nuisance variables (six motion parameters, whole-brain signal, ventricular signal, and white matter signal) and their temporal derivatives (computed by backward differences) were derived from BOLD data. All signals were regressed out using 3dTproject (AFNI v19.1.05102,111). This was followed by bandpass filtering at 0.01–0.1 Hz using 3dBandpass (AFNI v19.1.05102,111). Age-matched MNI-space volumetric data were then spatially smoothed using a 3 mm full width at half-maximum (FWHM) kernel. The decision to use a 3 mm kernel was based on our motivation to smooth the infant data less than the adult data, due to infants’ smaller brain size, compared with previous adult precision mapping that used a 4 mm kernel.

### Head movement and data quality in the medial temporal lobe

Head motion is known to dramatically affect functional connectivity estimates and to introduce spurious correlation patterns. This challenge is particularly pronounced in infant imaging. To minimize the effect of head motion on our connectivity findings, all BOLD runs with a maximum absolute head displacement greater than 2.2 mm were discarded from the analysis. Additionally, imaging the brain regions in the anterior MTL is a longstanding challenge since these regions are particularly prone to signal loss and distortion due to magnetic inhomogeneities seen clearly in T2∗-weighted images (BOLD data). To estimate the overall data quality, we calculated tSNR in every voxel by averaging the signal intensity in all available BOLD runs (prior to nuisance variables regression, filtering and smoothing) for each participant and dividing it by the standard deviation over time. The tSNR maps were averaged separately for each participant to provide a single tSNR value for each voxel in the brain.

### Discovery analysis

For studying the functional architecture of the cortical networks that are known to support episodic memory in adults and their associations with the infant hippocampal-entorhinal system, we adopted a seed-based discovery and validation framework^24^. In this approach, data for each subject are divided into two separate datasets – the discovery dataset and the validation dataset. The discovery dataset is used for generating hypotheses pertaining, for example, to network topography; the validation dataset is used for independent replication of the hypotheses generated with the discovery dataset in each individual subject. To this end, for each infant we divided all available BOLD data into discovery and validation datasets. All odd-numbered runs were assigned to a discovery dataset and all even-numbered runs were assigned to a validation dataset. After removing BOLD runs due to excessive head displacement, we had a range of 9 (54 min; 1 subject) to 12 (72 min; 2 subjects) BOLD runs for each individual subject. Previous precision mapping studies in adults suggest that approximately 30 min of BOLD data for each individual yields Pearson correlation coefficients > 0.9 with the “ground truth” individualized connectome. Therefore, we reasoned that having approximately 30 min of BOLD data for the discovery and validation datasets for each infant subject would result in a robust examination of distributed connectivity patterns.

Since there is no consensus pertaining to the cortical network organization of the infant brain, we adopted a seed-based and anatomically-driven exploration of the distributed circuitry in each individual infant. We used known adult individual-level distributed networks’ topography as a general reference; these findings are highly robust and most importantly, they have a strong biological basis, such that many of these networks can be identified also in adult macaques. All connectivity analyses were performed on volumetric data in age-matched MNI-space since we were interested in using a common reference frame. Seed regions in the study were individual voxels placed in various cortical regions (see below). For every seed used in the study, we calculated whole-brain Pearson correlation coefficients between the seed voxel’s time series and all other voxels’ time series for every discovery BOLD run using the instacorr function in AFNI. Resulting Pearson correlation coefficients were Fisher z-transformed and averaged across all discovery dataset BOLD runs. Functional connectivity maps for every seed voxel were thresholded using a spurious correlation threshold of z(r) > 0.15^15^. Negative correlation values, as well as correlation values in the white matter and ventricles were not interpreted.

The discovery analysis in each infant started with positioning a seed in the parahippocampal cortex 7 mm anteriorly to the last slice with visible tail of the hippocampus. The choice to begin the discovery analysis with the parahippocampal cortex was motivated by previous precision mapping studies showing that the cortical network DN-A in adults is exclusively associated with the parahippocampal cortex. We decided to place the seed 7 mm anteriorly from the tail of the hippocampus in order to remove the seed from the lingual gyrus and the retrosplenial cortex, to avoid network blurring. On the other hand, we avoided moving the seed too anteriorly, to avoid sampling signal from the entorhinal cortex. Anatomical landmarks were identified in each infant using an anatomical atlas containing adult human brain photos, the corresponding structural MR images and detailed anatomical labels at different slices^113^.

Seeds placed in the infant parahippocampal cortex resulted in a whole-brain distributed topography that strongly resembled adult cortical network DN-A (not shown). Next, we identified a region in the retrosplenial cortex (defined as the ventral part of the posterior midline cortex), which showed maximum Fisher z-transformed Pearson correlation coefficients with the parahippocampal seed and used this region as the defining seed of the hypothesized cortical network DN-A (see the upper panel in Figure 2 for hypothesized DN-A in two example participants; see Figure S2 for hypothesized DN-A in other participants). In this study, we specifically focused on the posterior midline cortex for identifying seeds for hypothesized cortical networks DN-A, DN-B and PMN for three reasons. First, evidence from neonatal rats points to adult-like connectivity of the hippocampal system with the retrosplenial cortex^41^. Second, previous adult precision mapping studies indicate that cortical networks DN-A, DN-B and PMN/SAL are well known to occupy anatomically adjacent regions in the posterior midline cortex – the retrosplenial cortex, caudal part of the posterior cingulate cortex and rostral part of the posterior cingulate cortex, respectively. Therefore, focusing on the posterior midline cortex would dramatically reduce the anatomical degrees of freedom in identifying seed regions associated with each cortical network. Third, since we were interested in examining the associations of these networks with the hippocampal-entorhinal system, our aim was to sample seeds that are located approximately equally distant from the hippocampal-entorhinal system, especially given recent evidence that long-range connectivity in the infant brain is weaker than short range connectivity^42–44^.

Next, using the topography of the putative DN-A as an anatomical anchor and based on previous precision mapping studying showing that the adult cortical networks DN-B and PMN/SAL associate with distinct parts of the posterior cingulate cortex, we explored the posterior midline cortex dorsal to the retrosplenial seed (putative DN-A). Our goal was to identify seed regions that were associated with distributed whole-brain topographies, which contain known adult DN-B and PMN/SAL components. These voxels were identified by visualizing the connectivity maps at a threshold of z(r) > 0.15 (above the noise correlation level) and by inspecting the anatomical locations of different cortical areas that consistently correlated with the posterior cingulate seeds across all infants - anterior midline, parietal lobe, lateral frontal cortex, lateral surface of the temporal lobe, intraparietal sulcus, precuneus, hippocampus and entorhinal cortex. It is important to emphasize that even though the explored seed regions maximized potential networks separation, these seeds and their associated distributed brain regions represented only hypotheses for distinct cortical networks and their associations with the hippocampus and the entorhinal cortex. Overall, the discovery datasets produced consistent hypotheses for the existence of distinct cortical networks. Therefore, a critical step was to put these hypotheses to an independent test and to examine whether distributed connectivity patterns observed using the discovery datasets will be replicated using independent validation datasets.

### Validation analysis

After identifying the seed regions in the posterior midline that maximized separation in the discovery datasets between putative networks DN-A, DN-B and PMN/SAL, we defined distributed network dissociation regions as voxels that showed maximum Fisher z-transformed Pearson correlation coefficients with the posterior midline seeds. These dissociation regions were defined *a priori* using only the discovery datasets. The distributed dissociation regions were positioned in the parietal lobe, dorsolateral frontal cortex, ventromedial prefrontal cortex, lateral surface of the temporal lobe, precuneus, intraparietal sulcus, parahippocampal cortex, anterior and posterior hippocampus, and different parts of the entorhinal cortex medial-lateral axis. Where possible, dissociation regions were always identified in the hemisphere ipsilateral to each posterior midline seed. Since the validation datasets were analyzed blindly to the discovery datasets, the critical test was to see whether the a priori chosen seed regions in the posterior midline cortex and the distributed dissociation regions will replicate the preferred connectivity patterns observed during the discovery analysis. For testing the hypothesis that distinct posterior midline seeds are associated with distinct cortical networks as well as with distinct parts of the hippocampal-entorhinal system, the connectivity patterns derived from the validation datasets were statistically tested using F-tests with one-way or two-way repeated measures analysis of variance (ANOVA). One-way ANOVA was used for testing the dissociations in the precuneus, intraparietal sulcus, parahippocampal cortex and lateral surface of the temporal lobe. Two-way ANOVA was used for testing the dissociations in the inferior parietal lobule, dorsolateral frontal cortex, ventromedial prefrontal cortex, anterior and posterior hippocampus, and entorhinal cortex.

In addition to the formal statistical testing, we examined “model-free” whole-brain associations with distinct parts of the entorhinal cortex medial-lateral axis. We selected entorhinal seeds that showed maximum network separation between DN-A and PMN/SAL in the discovery analyses and calculated whole-brain connectivity patterns of these seeds using validation datasets. The aim of this analysis was to see whether differential associations with the hippocampal long-axis, as well as network topographies revealed using discovery datasets will be replicated using “back-projections” connectivity using validation datasets.

### Entorhinal cortex mask

We used previous anatomical studies in human adults to define the entorhinal cortex in each individual infant^114,115^. Although we acknowledge the limitation in using adult-based procedures for defining the infant entorhinal cortex, to the best of our knowledge, there is no existing work examining the anatomical properties, as seen using MRI, of the entorhinal cortex in the human infant brain. Using MR structural data, we localized major anatomical landmarks, such as the amygdala, collateral sulcus, and the ambient gyrus, for defining the entorhinal cortex. An adult atlas presenting postmortem brain slices and the corresponding MR images was used as an anatomical reference^113^. The anterior boundary of the entorhinal cortex was defined 3 mm anterior to the first coronal slice with visible hippocampus, at the level of the amygdala and the anterior part of the ambient gyrus, which is part of the entorhinal cortex based on human cytoarchitectonic data^116^. The posterior boundary of the entorhinal cortex was defined by the end of the uncus. On the medial-lateral axis, the medial border of the entorhinal cortex was defined by the parahippocampal gyrus, likely resulting in inclusion of parts of the pre/parasubiculum. As the lateral border of the entorhinal cortex, the shoulder of the collateral sulcus was used.

In order to formally examine changes in networks’ associations with the infant entorhinal cortex, we defined for each infant a bilateral mask comprising the presumed intermediate entorhinal cortex band. The mask was defined bilaterally along the entorhinal anterior-posterior axis and between the local DN-A and PMN/SAL maximum correlation coefficient on each coronal slice on the medial-lateral axis. The changes in percentage of noise-passing voxels across time point one and time point two were tested using a two proportions z-test.

### Longitudinal changes of cortical network associations with the infant medial temporal lobe

To study how associations of the cortical networks DN-A, DN-B and PMN/SAL with the hippocampal-entorhinal system change over time, we invited the parents of the 5 original infants for an additional session approximately 9 months after the last scanning session. While all parents returned for the additional scanning session, we managed to collect longitudinal MRI data only from three infants (other infants did not remain asleep in the scanner bore) and data from the infants for whom we collected more than 20 min of high-quality fMRI data (Table 1) were analyzed for the purpose of this study. After removing BOLD runs due to excessive head displacement, we retained 4 BOLD runs (24 min) of additional data for P1 and 8 BOLD runs (48 min) of additional data for P2 at time point 2. The goal of the analysis was to examine potential developmental changes in the association of the cortical networks DN-B, PMN and DN-A with the infant medial temporal lobe. To this end, we projected the discovery seeds for each network and for each infant’s brain at time point 1 to the same infant brain at time point 2. We then examined associations of these projected seeds in the hippocampal-entorhinal system. In order to formally examine change in networks’ associations with the infant hippocampus, for each network we calculated the percentage of the hippocampal voxels (using a bilateral mask) that passed the noise correlation threshold (z(r)>0.15) at both time points.

### Replication analysis using parietal seeds

To demonstrate that the dissociation between DN-A and DN-B in the neocortex does not depend on specific seed selection in the posterior midline cortex, we performed a replication of validation analysis by placing new seeds in the inferior parietal cortex using discovery datasets in each participant. Parietal seeds were defined as neocortical regions in the parietal cortex that showed local maximum Fisher z-transformed correlation coefficients with the original posterior midline seeds used to define the cortical networks DN-A and DN-B. See Figure S11 for cortical networks DN-A and DN-B defined using parietal seeds. This analysis replicated the reported dissociated between DN-A and DN-B reposted in Figure 4, except for a not significant interaction in the ventromedial prefrontal cortex in P2 and a significant interaction in the ventromedial prefrontal cortex in P1 and P2 that did not show a cross-over effect.

### Assessment of infants’ general development

Infants’ general development was measured using the standard Bayley Scale of Infant and Toddler Development, Third Edition (Bayley-III)^40^. Infants performed multiple tasks targeting various aspects of cognitive and motor development, including language and fine motor skills. All behavioral testing was performed prior to MRI testing. Raw Bayley Scale scores were used for each infant. Additionally, at 24 months of age P1 and P2 performed various episodic memory tasks that were not analyzed.

