## Supplementary Figures S1-S10 for "Subregions of the infant hippocampal-entorhinal system are linked to distinct cortical networks in the first two years of human life"

### Supplementary Materials

Figs. S1 to S10

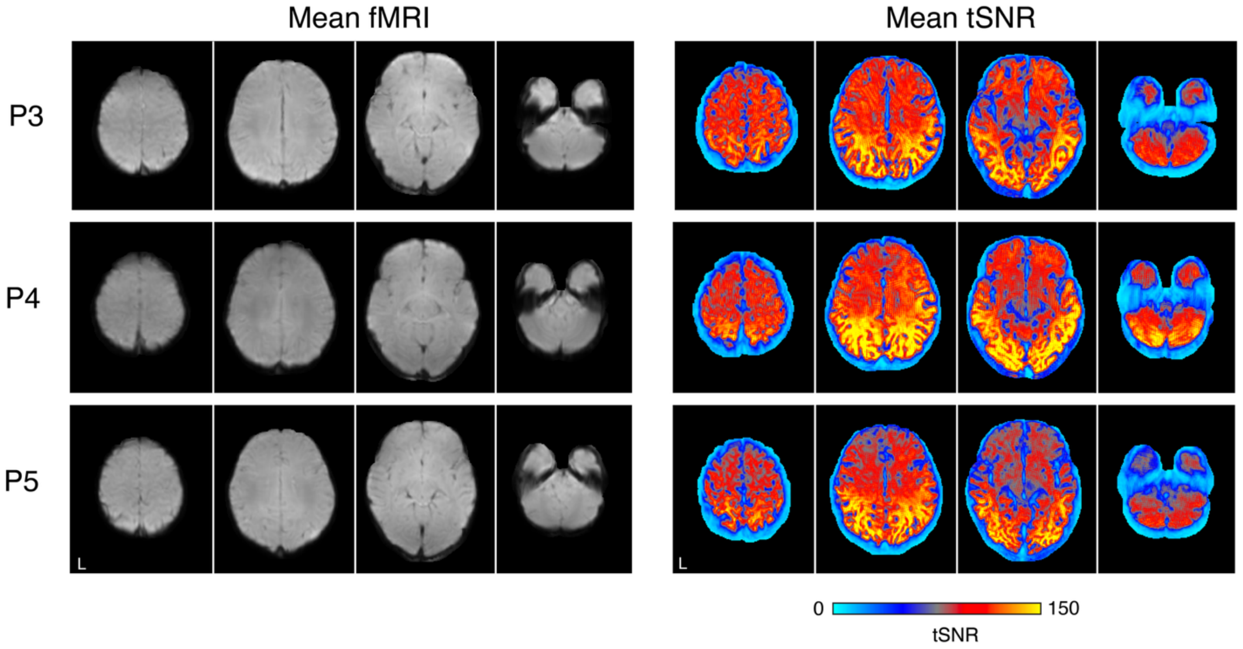


**Figure S1. Infant fMRI data quality.** Upper row – whole brain data quality. Axial slices from S3, S4 and S5 show mean BOLD (left) and mean tSNR (right) data. High coverage and high tSNR were obtained in most regions of the brain, particularly in the medial temporal lobe.

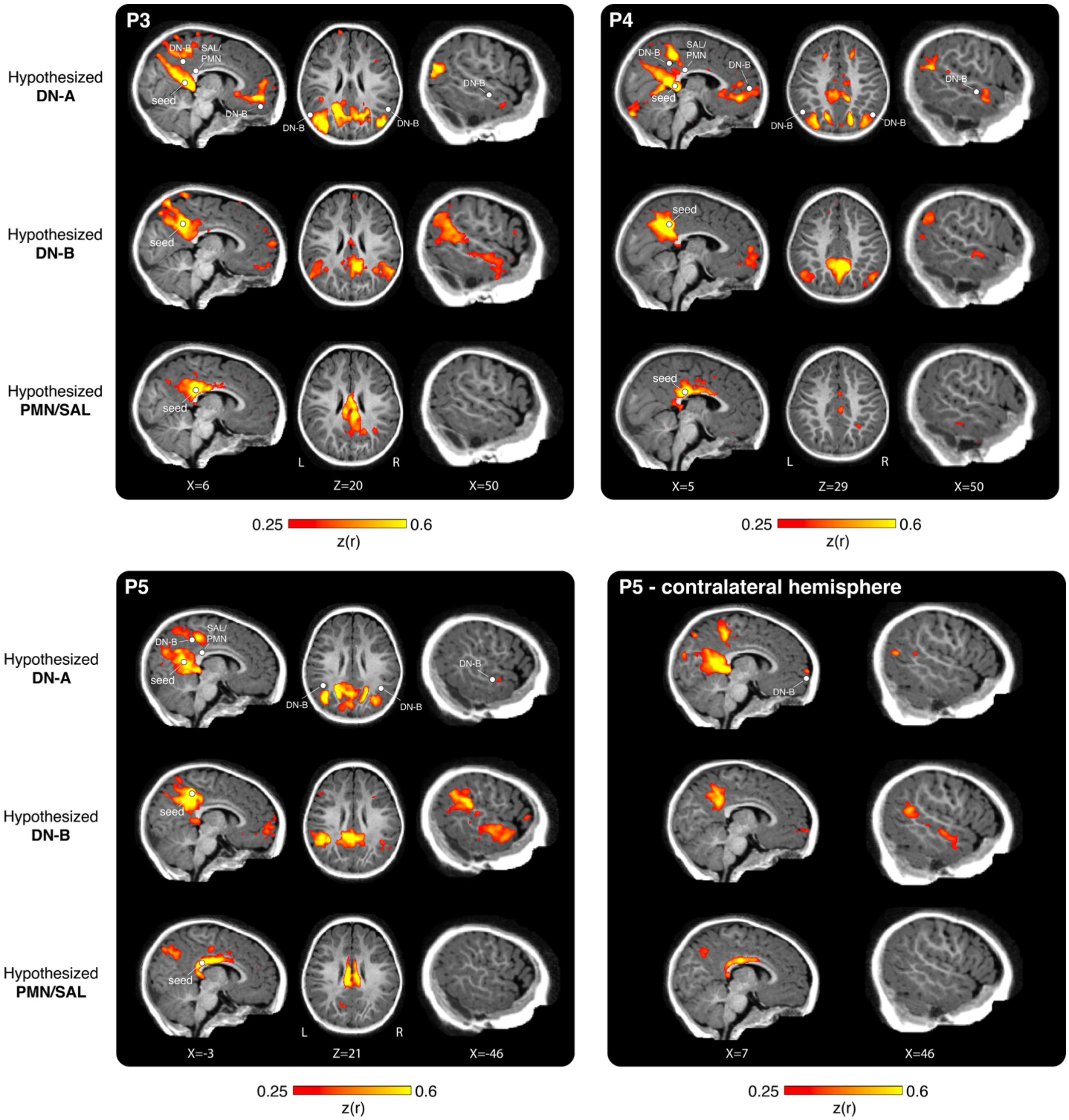
P3
P4
Hypothesized DN-A
Hypothesized DN-B
Hypothesized PMN/SAL
DN-B
SAL/ PMN
seed
L
R
X=6
Z=20
X=50
X=5
Z=29
X=50
0.25
0.6
z(r)
P5
P5 - contralateral hemisphere
Hypothesized DN-A
Hypothesized DN-B
Hypothesized PMN/SAL
X=-3
Z=21
X=-46
X=7
X=46

**Figure S2. Network discovery.** Using seeds placed in the posterior midline cortex we identified candidate networks DN-A (top row), DN-B (middle row) and PMN/SAL (bottom row) in the infant brain in P3, P4 and P5. Note the close anatomical position of the DN-A, DN-B and PMN/SAL seeds indicated by white circles in the top row for each participant. All three networks resembled their adult counterparts and occupied anatomically adjacent brain regions indicated by white circles in the parietal lobe, ventral medial prefrontal cortex, lateral prefrontal cortex and lateral surface of the temporal lobe. Regions indicated by the circles were defined as the local maximum Fisher transformed Pearson correlation coefficient with the posterior midline seeds for each network in each presented brain slice. These regions were then submitted for an independent validation test presented in Figure 4 and Figure 5. See Figure 2 for data from other participants and Figure S3 for group-level surface maps.

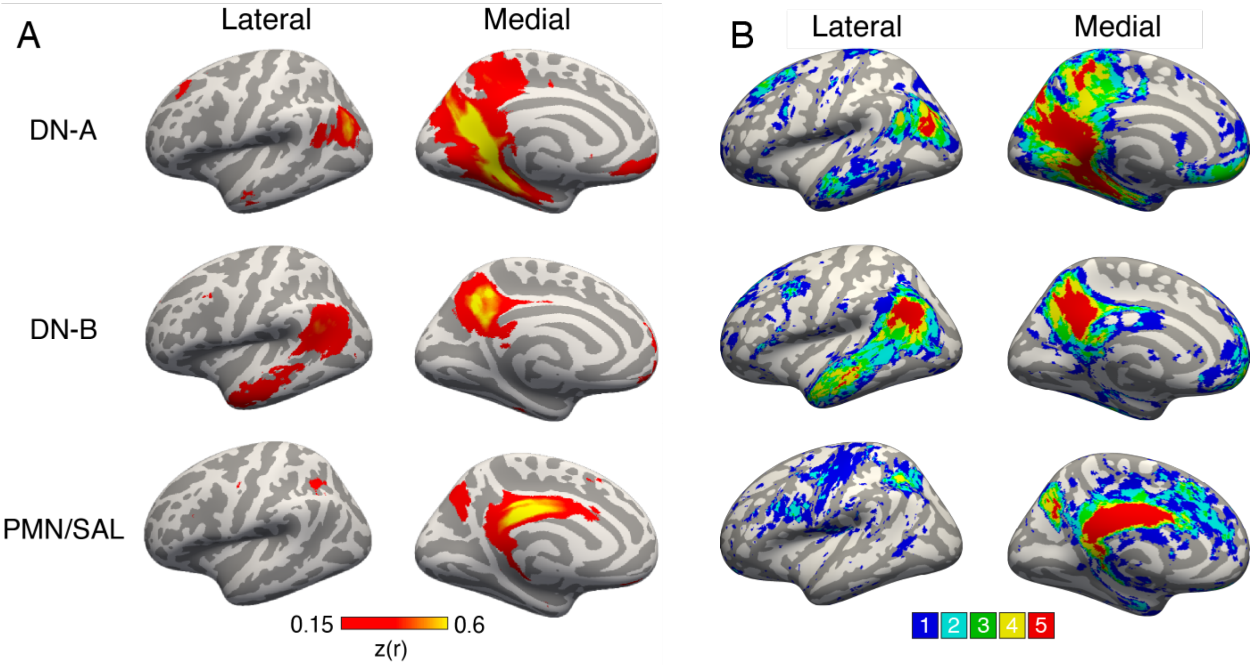


**Figure S3. Group-average cortical surface representations of the discovery datasets and group overlap maps.**

**(A)** We projected mid-cortical volumetric connectivity values from the discovery dataset of each individual to a common fsaverage5 cortical surface of the left hemisphere in order to generate group-level, average cortical surface representations of connectivity measures for the cortical networks DN-A (top), DN-B (middle), and PMN/SAL (bottom). See Figure2 and Figure S2 for volumetric maps of individual participants.

**(B)** For each participant, we overplayed binarized connectivity maps (using a threshold of z(r)>0.15) on a common fsaverage5 cortical surface of the left hemisphere in order to estimate the amount of spatial convergence across individuals for the cortical networks DN-A (top), DN-B (middle), and PMN/SAL (bottom).

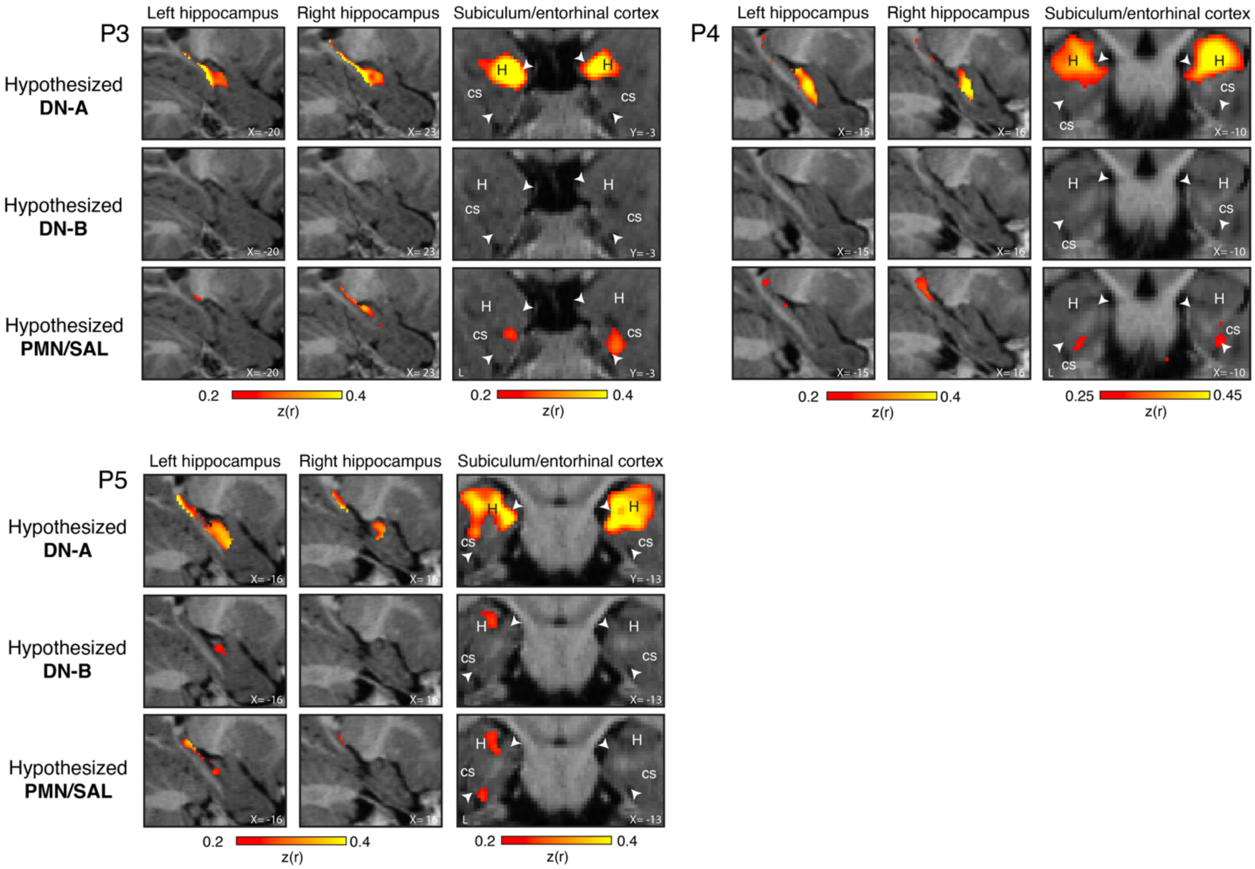


**Figure S4. Network discovery in the medial temporal lobe.** Using seeds placed in the posterior midline cortex of P3, P4 and P5 we examined candidate networks DN-A (top row), DN-B (middle row) and PMN/SAL (bottom row) associations with the infant brain medial temporal lobe structures, specifically the hippocampus and the entorhinal cortex. The infant cortical network DN-A was associated with the anterior hippocampus and the most medial part of the entorhinal cortex, similar to adults. The infant cortical network PMN/SAL was associated with the posterior hippocampus and the most lateral part of the entorhinal cortex, similar to adults. Surprisingly, associations of the infant DN-B with the medial temporal lobe structures were minimal and not consistent. See Figure 3 for data from other participants.

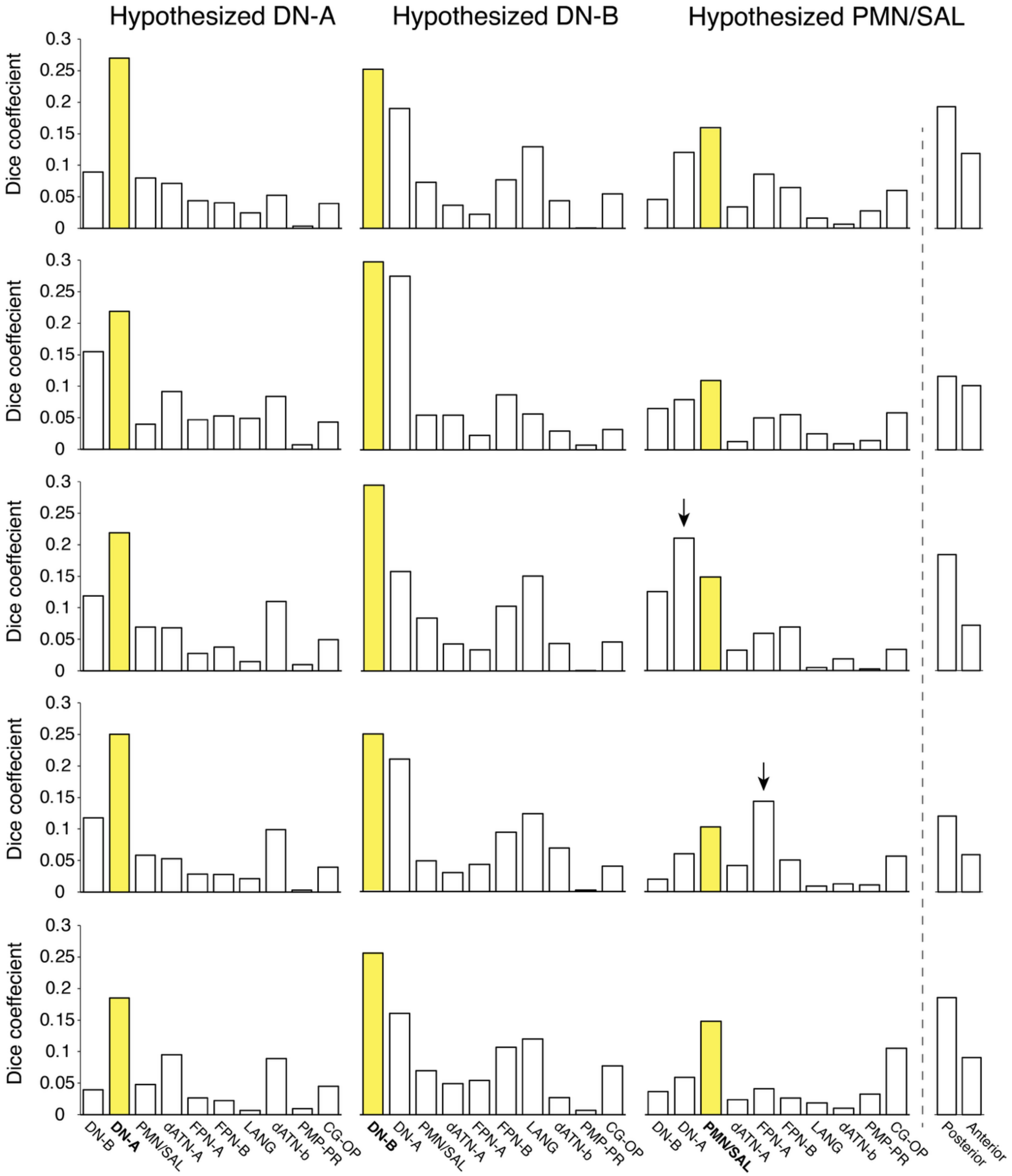

Hypothesized DN-A
Hypothesized DN-B
Hypothesized PMN/SAL
Dice coeffecient
0
0.05
0.1
0.15
0.2
0.25
0.3
DN-B
DN-A
PMN/SAL
dATN-A
FPN-A
FPN-B
LANG
dATN-b
PMP-PR
CG-OP
Posterior
Anterior

**Figure S5. Dice coefficient analysis.** We calculated dice coefficients between binarized connectivity maps (thresholded at $z(r)>0.15$) for each cortical network of interest (DN-A, DN-B, and PMN/SAL) and their adult counterpart in the DU15 15-network parcellation. We found that the networks that we identified using seed-based analysis had the highest dice coefficients with their parallel networks in the adult brain, except for PMN/SAL in P3 and P4 (marked with arrows). Furthermore, to quantify the missing frontal components of the infant PMN/SAL, we calculated dice coefficients separately for the anterior and posterior parts of the infant and adult PMN/SAL. As can be seen in the right column, in all participants, the dice coefficient was greater for the posterior component compared with the anterior component.

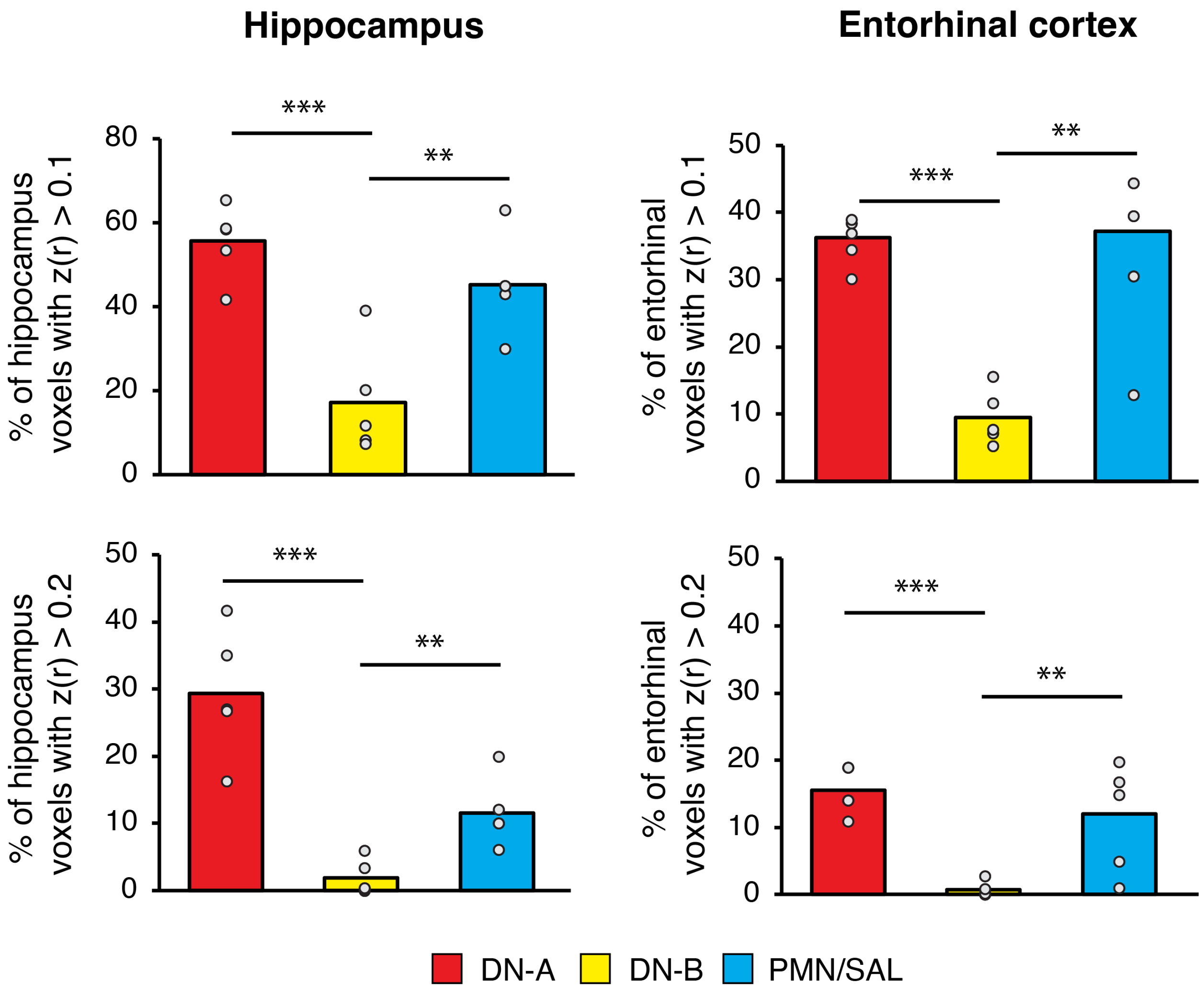


**Figure S6. Validation of weaker DN-B associations with the infant medial temporal lobe using different noise thresholds.** Using independent validation datasets, we examined the average percentage of threshold-passing voxels using a threshold of z(r)>0.1 (top) and z(r)>0.2 (bottom) in the hippocampus and the entorhinal cortex, associated with DN-A, DN-B and PMN/SAL. All brain regions used for this validation analysis were defined a priori using discovery datasets. Associations with the cortical network DN-B were significantly lower than with DN-A and PMN/SAL (all p-values < 0.01). Bars represent mean percentage of threshold-passing voxels across all seeds and participants, presented as individual dots. Numbers in parentheses represent the age of each infant, m – months. $**p < 0.01$, $***p < 0.001$. See Figure 6 for the same analysis using a noise-level threshold of z(r)>0.15.

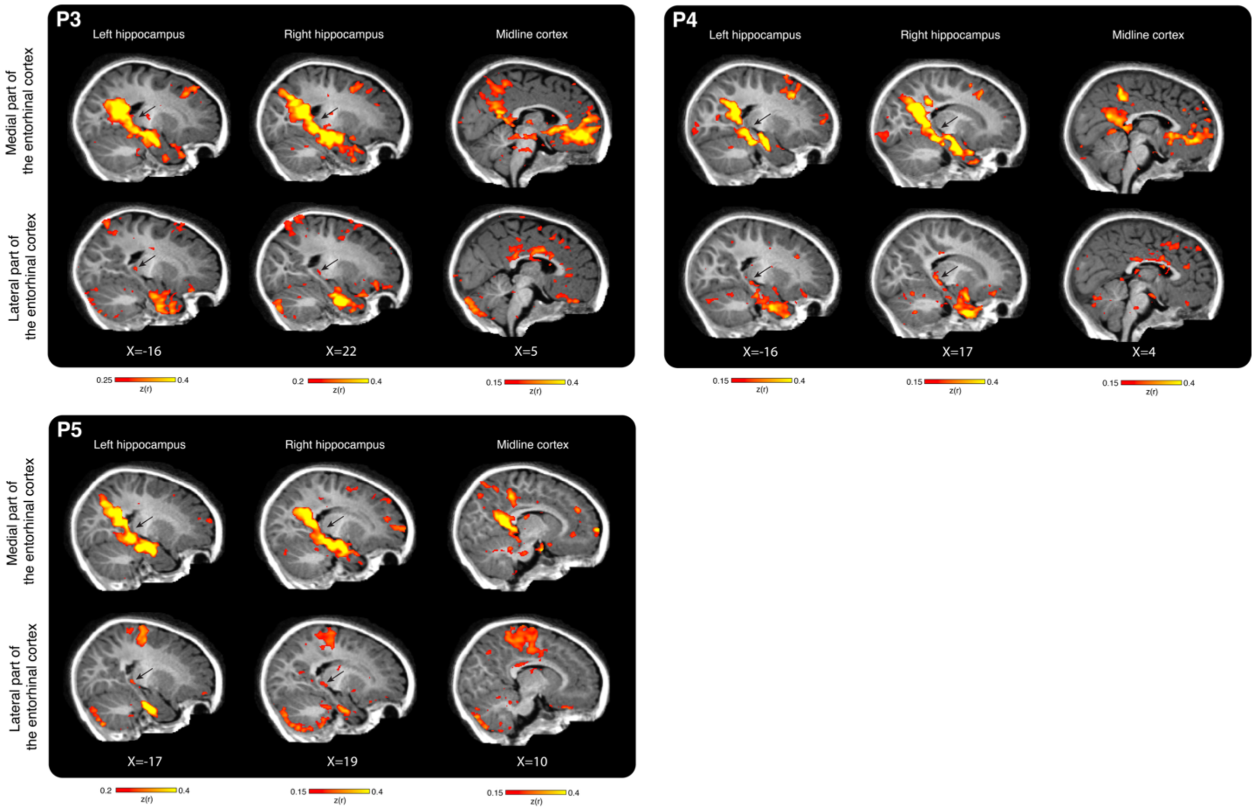


**Figure S7. "Back projections" from the entorhinal cortex.** In addition to formal statistical testing, we examined whether seeds placed in distinct parts of the infant entorhinal cortex, informed by the discovery dataset, would associate with distinct parts of the hippocampal long axis and distinct cortical networks in the validation dataset. We found that seeds placed in the most medial and lateral parts of the infant entorhinal cortex revealed whole brain topographies corresponding to infants' DN-A and partially to PMN/SAL, including anterior and posterior hippocampus (marked with black arrows) respectively. These results provide further support for the existence of distinct entorhinal-hippocampal circuits in the infant brain that associate distinct parts of the entorhinal cortex with distinct parts of the hippocampus. See Figure 7 for data from other participants.

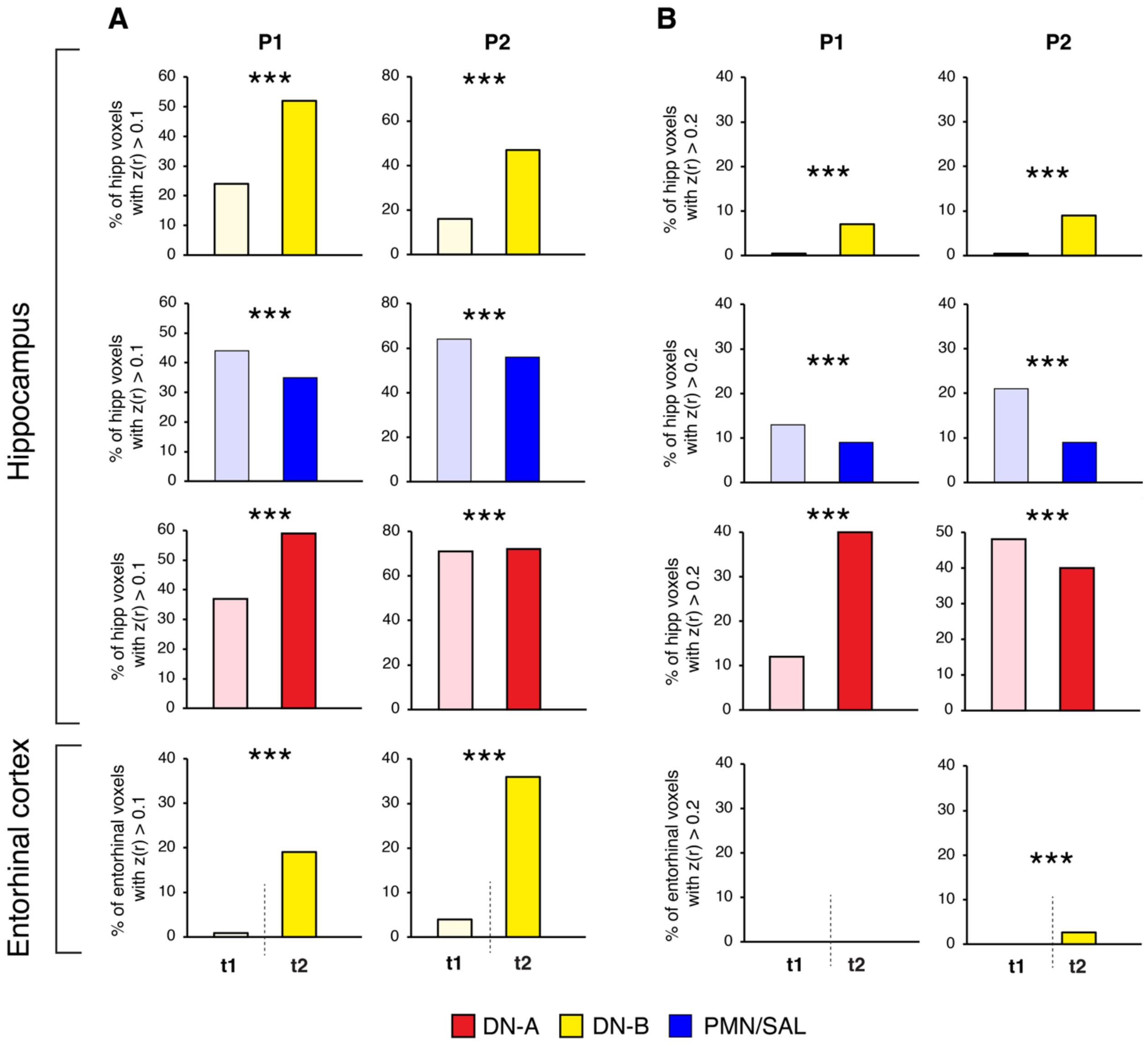


**Figure S8. Exploratory analysis examining longitudinal changes in cortical network associations with the infant medial temporal lobe using different noise thresholds.** We performed the same analysis presented in the main Figure 8 examining changes in cortical networks' associations with the hippocampal-entorhinal system between 15 (t1) and 24 (t2) months of age. All results reported in the main Figure 8 we replicated using both (A) a more lenient threshold of z(r)>0.1, and (B) a more conservative threshold of z(r)>0.2. Note that in P1, no entorhinal voxels passed the conservative threshold in neither time point. See Figure 8 for the same analyses using a noise threshold of z(r)>0.15; ** p< 0.01, *** p<0.001.

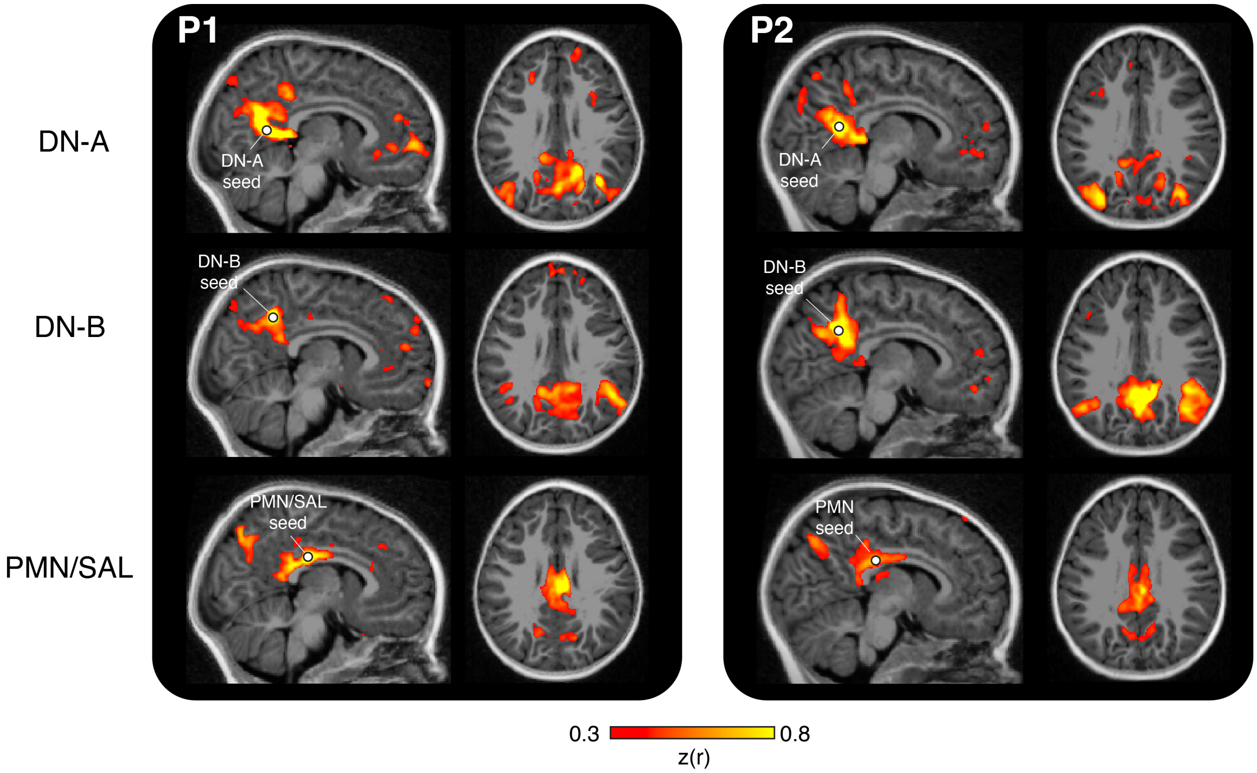


**Figure S9. Cortical networks at time point 2.** Using the same seeds in the posterior midline cortex as in time point 1, we identified cortical networks DN-A (upper row), DN-B (middle tow) and PMN/SAL (bottom row) in P1 and P2 that were nearly 2 years of age. Compare to similar networks' topographies of the same participants presented in Figure 2 when the same infants were approximately 8 months younger.

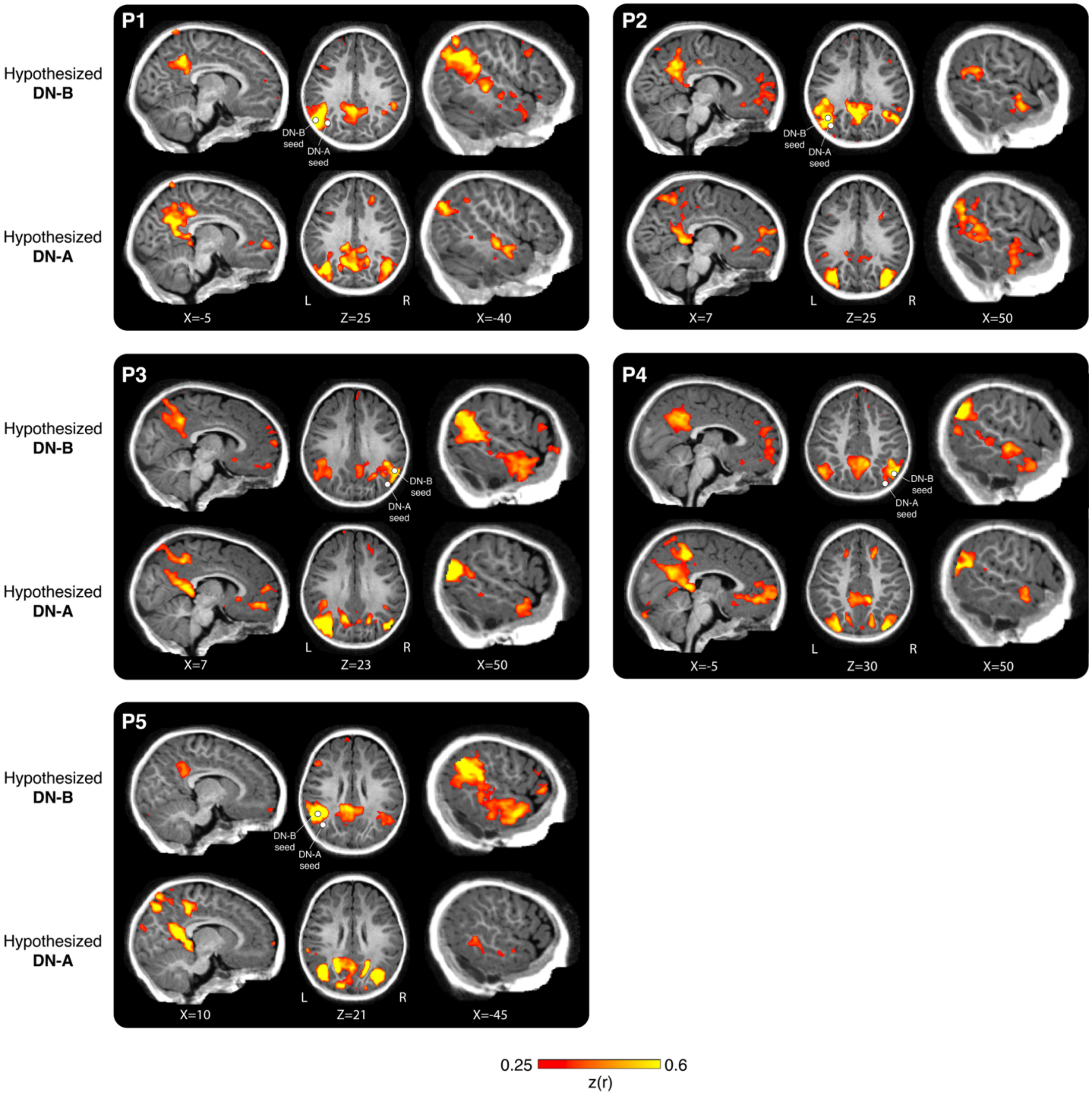


**Figure S10. Cortical networks DN-A and DN-B defined using parietal seeds.** Using discovery datasets, we defined new seeds located in the parietal cortex that showed the strongest connectivity with the original seeds placed in the posterior midline cortex. Using the new parietal seeds, new dissociation points in the lateral surface of the temporal lobe, ventromedial prefrontal cortex and dorsolateral prefrontal cortex were defined (posterior midline seeds remained unchanged). All reported results dissociating cortical networks DN-A and DN-B presented in Figure 4 were replicated, except for a not significant interaction in the ventromedial prefrontal cortex in P2 and a significant interaction in the ventromedial prefrontal cortex in P1 that did not show a cross-over effect.
